# Multi-species integration, alignment and annotation of single-cell RNA-seq data with CAMEX

**DOI:** 10.1101/2025.01.25.634864

**Authors:** Zhen-Hao Guo, De-Shuang Huang, Shihua Zhang

## Abstract

Single-cell RNA-seq (scRNA-seq) data from multiple species present remarkable opportunities to explore cellular origins and evolution. However, integrating and annotating scRNA-seq data across different species remains challenging due to the variations in sequencing techniques, ambiguity of homologous relationships, and limited biological knowledge. To tackle the above challenges, we introduce CAMEX, a heterogeneous Graph Neural Network (GNN) tool that leverages many-to-many homologous relationships for multi-species integration, alignment, and annotation of scRNA-seq data from multiple species. Notably, CAMEX outperforms state-of-the-art methods integration on various cross-species benchmarking datasets (ranging from one to eleven species). Besides, CAMEX facilitates the alignment of diverse species across different developmental stages, significantly enhancing our understanding of organ and organism origins. Furthermore, CAMEX enables the detection of species-specific cell types and marker genes through cell and gene embedding. In short, CAMEX holds the potential to provide invaluable insights into how evolutionary forces operate across different species at single-cell resolution.

## Introduction

Single-cell omics techniques enable the profiling of thousands of cells in one experiment, revealing a comprehensive landscape of heterogeneous cells [1]. These techniques facilitate the discovery of rare cell states, infer gene regulatory networks, and enhance our understanding of development and differentiation processes at a finer resolution [2]. With advancements in sequencing methods, scRNA-seq data from different species, including humans [3], monkeys [4], mice [5], zebrafish [6], and more, have been generated by various studies. These datasets provide clues on the origin of cellular diversity and evolutionary drivers of cellular morphology [7]. Diverse studies have revealed many cryptic cell types across multiple species [8]. Comparing gene expression of individual cells across species uncovers species-specific transcriptomic regulation [9], provides novel insights into the phylogeny of cells and organs, and elucidates how cells function in different developmental stages [10]. Integrating scRNA-seq data across species advances our understanding of molecular and cellular biology and becomes an essential first step toward comparative biology [11, 12]. These scRNA-seq data enable cross-species comparisons and analyses, shedding light on cellular functions and evolutionary dynamics. Only through these comparisons can we eventually decipher and reconstruct species evolution [13].

Single-cell RNA sequencing experiments can only process a limited number of cells every time. Data from various batches exhibit significant variations due to differences in capture time, operating personnel, and sequencing platforms [14]. These batch effects can obscure biological signals and interfere with our ability to analyze specific biological changes of interest [15]. Researchers have developed many methods, such as Harmony [16], scVI [17], and SingleCellNet (SCN) [18], for integrating and annotating scRNA-seq data within a single species. However, existing methods align features in the expression profile matrix across different species solely through one-to-one homologous genes, resulting in the loss of much biological information [19]. Several methods have made initial explorations into the integration and annotation of cross-species scRNA-seq data, such as BPA (Biological Process Activity) [20], SATURN [21] and CAME [22]. However, the limited biological knowledge available for non-model organisms poses challenges for implementing BPA. SATURN employs a protein-language model to measure gene similarity across different species; however, discrepancies can arise between proteome and transcriptome due to processes like alternative splicing.Our previous annotation method, CAME, leverages many-to-many homologous gene relationships but is limited to mapping a pair of species into the same embedding space.

In a word, integrating scRNA-seq data obtained from diverse sources and non-model organisms faces challenges due to technical and biological factors. First, the data from different sequencing platforms vary greatly, including the difference of detected UMIs and genes. Second, aligning the features of expression profile matrices from other species through the relationships between orthologous genes poses a significant difficulty. Last, processing scRNA-seq data from non-model organisms with limited biological knowledge, e.g., specific regulatory networks, distinctive communication mechanisms, and individual diversity, is a complex challenge.

To this end, we introduce CAMEX, a heterogeneous graph neural network (GNN) tool that leverages many-to-many homologous relationships for integration, alignment, and annotation of scRNA-seq data from multiple species. Specifically, we construct a heterogeneous graph consisting of cells and genes based on expression matrices and many-to-many homology relationships. We design a heterogeneous graph neural network as an encoder to encode cells and genes from different species into the common embedding space. For the batch correction task, we design two decoders including a dot-product decoder to reconstruct the edges in the heterogeneous graph and a full-connected decoder to reconstruct the cell expression. For the cell annotation task, we feed the cell embedding into the graph attention network (GAT) for classification. CAMEX achieved competitive performance compared to SOTA methods in integrating scRNA-seq data in different scenarios including the liver (involving four species with two technologies), ovary (encompassing three species with rare populations) and pancreas (comprising three species with multiple batches). Furthermore, CAMEX successfully integrated a testis dataset across 11 different species, preserving the sperm differentiation trajectory. Additionally, CAMEX correctly integrated RNA-seq data from seven species, spanning multiple organs and various developmental stages. Notably, CAMEX could align and predict developmental time beyond the Carnegie stages across different species. Finally, we demonstrated that CAMEX effectively annotates multi-species cell labels, even for distantly related species, and can discover new subgroups and marker genes.

## Results

### Overview of CAMEX

CAMEX takes diverse single-cell datasets from various species generated using the same or different technologies as input (**Fig. 1**). The expression matrix can be transformed into a bipartite graph *A*, with cells and genes as vertices. An edge *A*_*nm*_ is established between a cell *n* and a gene *m* if the cell *n* expresses the gene *m*. Then, we construct a *k*-nearest neighbor (kNN) graph to connect cells according to their expression similarity. In addition, self-loops are added for each cell and gene to prevent excessive smoothing in the GNN’s message passing. Finally, we connect the genes of different species based on many-to-many homologous relationships. The initial feature of each cell node corresponds to its original expression, while the initial feature of each gene node is set to empty. CAMEX employs a heterogeneous GNN encoder to embed each node into a low-dimensional common space by nonlinearly propagating features from neighboring nodes to center nodes.

**Figure 1.**
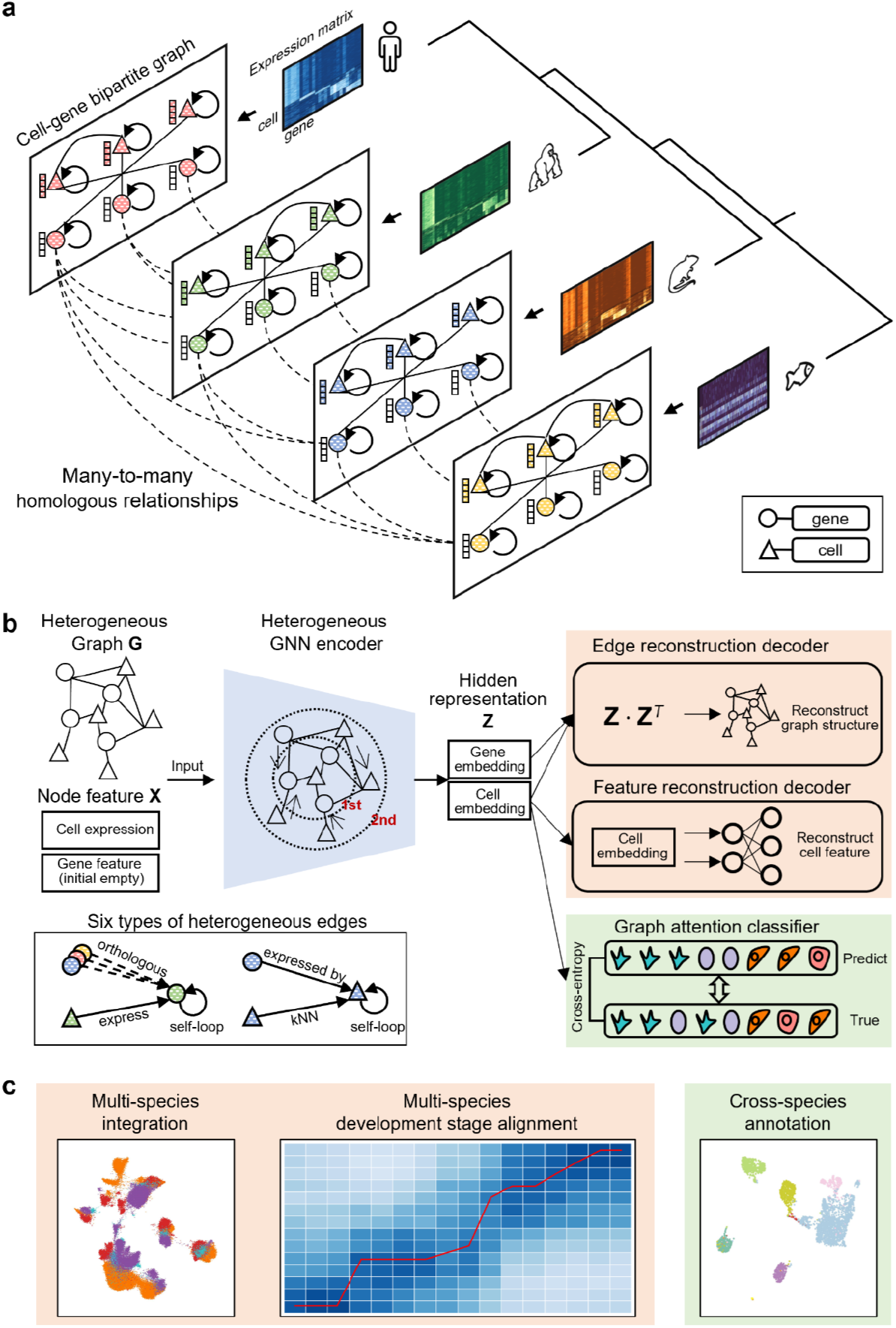
Overview of CAMEX. **a**, Construction of a heterogeneous graph representing the relationships between cells and genes across different species. This multi-species cell-gene heterogeneous graph consists of six types of edges, i.e., ‘cell express gene’, ‘gene expressed by cell’, ‘cell kNN’, ‘gene orthologous relationship’, ‘cell self-loop’, and ‘gene self-loop’. **b**, CAMEX leverages a heterogeneous GNN encoder to embed cells and genes from various species into a shared space. The integration task employs the reconstruction decoders for both edges and features. The cell annotation task utilizes a graph attention network classifier. Training details outline the message propagation paradigm for six types of edges and graph sampling techniques. **c**, CAMEX serves several applications, including multi-species integration, alignment of developmental stages, and cell annotation.

As to the integration task, CAMEX utilizes an encoder-decoder architecture. Specifically, it employs a heterogeneous GNN as an encoder, a dot product decoder to reconstruct the edges in the cell-gene heterogeneous graph, and a fully connected decoder to reconstruct the cell expression. The loss function comprises both cross-entropy loss between the predicted and raw labels of edges and mean square error (MSE) loss between the predicted and raw expression of cells. CAMEX minimizes them to optimize the model parameters by the back-propagation algorithm in an unsupervised manner.

As to the annotation task, CAMEX adopts an encoder-classifier architecture with at least one given reference dataset containing cell labels. It employs a heterogeneous GNN as an encoder and a GAT to predict cell labels. The loss function computes the cross-entropy loss between the predicted and ground truth labels of cell annotations in the reference dataset. CAMEX minimizes it to optimize the model parameters by the back-propagation algorithm in a semi-supervised way.

We collected various cross-species scRNA-seq datasets to benchmark CAMEX and other baseline methods. In the integration task, we applied CAMEX to the datasets and compared its performance against other SOTA methods, i.e., Harmony [16], PyLiger [23, 24], scVI [17], and Scanorama [25]. For CAMEX, we leveraged many-to-many homologous gene relationships to construct heterogeneous networks. In contrast, we employed one-to-one homologous genes to align expression matrices for the baseline methods. We conducted a quantitative evaluation of batch correction and biological variation conservation. Specifically, the batch correction score using metrics including kBET (k-nearest-neighbor Batch Effect Test) [26], PCR batch (Principal Component Regression of the batch covariate) [26], ASW batch (Average Silhouette Width across batch) [26], and graph connectivity [14]. The biological variation conservation score using metrics including ARI (Adjusted Rand Index) [27], NMI (Normalized Mutual Information) [28], Cell-type ASW [14], ISO label F1 (Isolated label F1) [14], and ISO label silhouette (Isolated label silhouette) [14]. Higher values for each metric indicate better performance (**Methods**). Finally, we computed an overall score as a 4:5 weighted average of the batch correction score and the biological variation conservation score to represent the overall model performance. Similar to the integration task, here we applied CAMEX to the datasets and compared its performance against other baseline methods, i.e., SCN [18], scANVI [29], and Cell BLAST [30]. We conducted ACC (accuracy), Prec (precision), and F1 (F1 score) for the quantitative evaluation of each baseline method.

### CAMEX achieves superior integration performance in different multi-species scenarios

We curated three multi-species scRNA-seq datasets, comprising the liver data (involving four species with two technologies), ovary data (encompassing three species with rare populations), and pancreas data (comprising three species with seven batches by six different technologies) to simulate a diverse range of real-world scenarios for benchmarking CAMEX against other baseline methods. In the liver dataset, the scRNA-seq data for humans, macaques, and mice originating from the same study were generated using the same sequencing platform [31]. However, the zebrafish scRNA-seq data were obtained from the zebrafish atlas [13], where the detected number of Unique Molecular Identifiers (UMIs) and genes were relatively low (**Supplementary Fig. S1a, b**). Each of the datasets contains distinct cell types unique to that respective species. While the human, monkey, and mouse datasets exhibit balanced cell types, the zebrafish dataset predominantly consists of hepatocytes (**Supplementary Fig. S1c**). These two factors significantly increased the complexity of data integration. In this context, CAMEX was the only method capable of accurately integrating scRNA-seq data across different species and achieving the highest overall score (**Fig. 2a-c**). It effectively clustered hepatocytes, fibroblasts, cholangio, endothelial cells, B plasma cells, and Kupffer cells (macrophages). Compared to other methods, CAMEX established more distinct cell population boundaries. Although scVI also aggregated similar cell types across different species, its large intra-class distances led to lower kBET and ASW batch scores, positioning it below CAMEX for the overall ranking. PyLiger performed well in integrating hepatocytes but cannot address other cell types well. Harmony excelled in integrating non-hepatocytes but failed when dealing with hepatocytes. The integration result of Scanorama was similar to Harmony, but its overall performance lagged behind Harmony. In addition, all methods successfully integrated liver-specific macrophages (macrophages and Kupffer cells). We examined the expression of typical marker genes in each cell type and verified the identities of macrophages and Kupffer cells (**Fig. 2d**). Although the genes receiving top attention from each cell type vary considerably across species, they exhibit high cell-type specificity.

**Figure 2.**
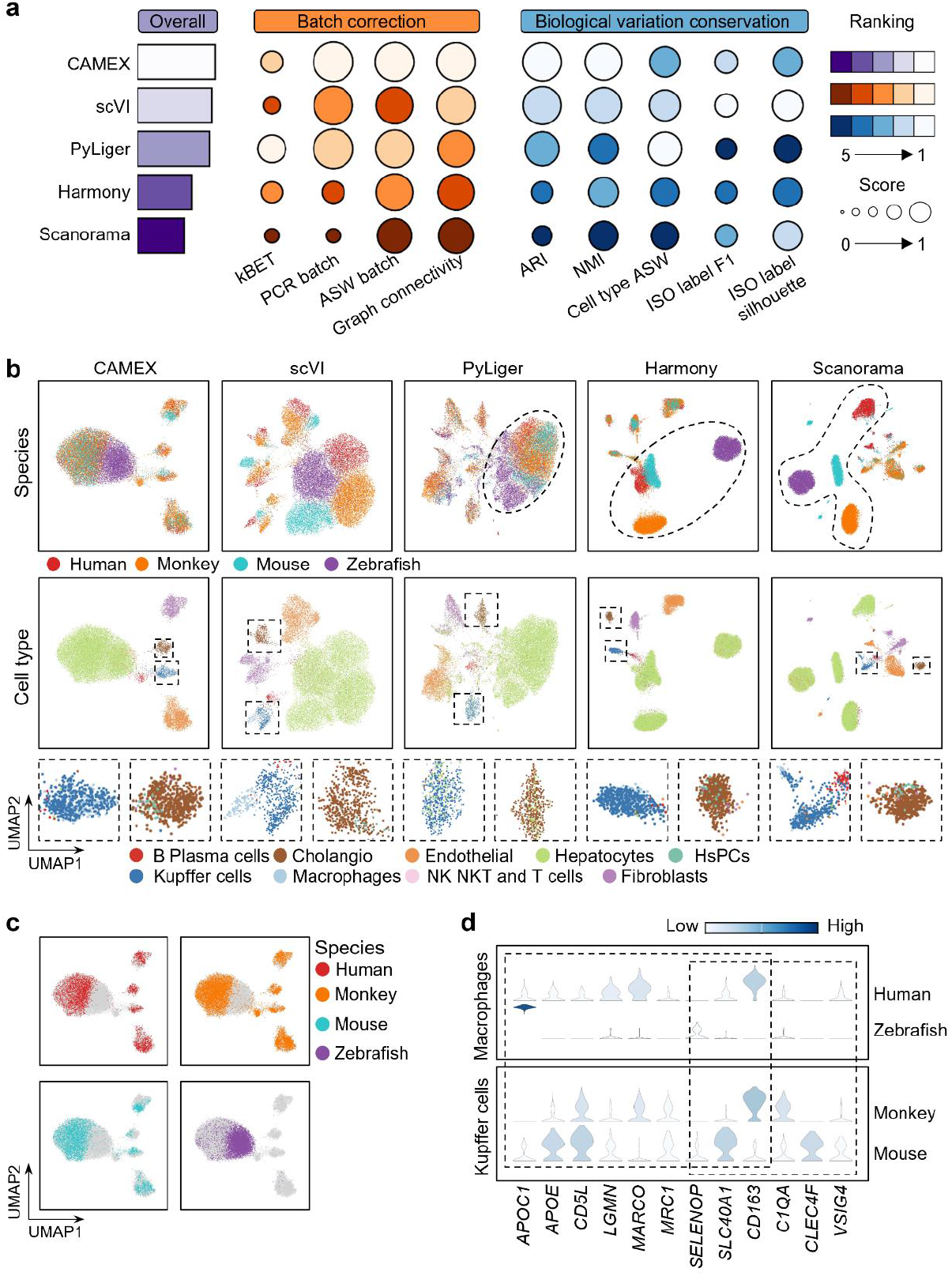
Benchmarking CAMEX and SOTA methods on the liver dataset across four species. **a**, Overall score evaluation on the liver. The overall score considers both batch correction and biological variation conservation performance. Rankings are depicted using a color gradient, where lighter colors indicate better performance. **b**, UMAP visualization of joint embedding space on the liver dataset. Colors represent species annotations (first row) and cell type annotations (second row). **c**, UMAP visualization of CAMEX integration result where color denotes different species. **d**, Marker genes of Macrophages and Kupffer cells in different species.

In the ovary dataset, the scRNA-seq data for humans [32-37], macaques [38], and mice [39] originating from different sequencing platforms resulted in distinct data types (**Supplementary Fig. S2a-c**). The macaque dataset contains a rare, unique oocyte cell type. In this scenario, CAMEX, PyLiger, and scVI can integrate the scRNA-seq data from three species while preserving this rare cell population, but CAMEX achieved a higher overall score (**Supplementary Fig. S2d-e**).

In the pancreas dataset, the scRNA-seq data came from humans [34], macaques [40], and mice [34] with humans containing five batches (by five different technologies) of data (**Supplementary Fig. S3a**). We used this to simulate a multi-species, multi-batch scenario. The human batches in this dataset contain roughly the same cell types. In addition, mice and macaques contain cell types that are subsets of humans (**Supplementary Fig. S3b, c)**. In this scenario, CAMEX can preserve the strong biological signal and effectively separate different cell types (**Supplementary Fig. S3d, e)**. Almost all methods, excluding Scanorama, exhibited relatively decent integration performance and can clearly distinguish various cell types. However, Harmony tended to over-integrate the acinar and ductal cell types. PyLiger over-integrated the acinar and ductal cells and the delta and gamma cells but failed to integrate the alpha and beta cell types. Conversely, scVI cannot distinguish the acinar and ductal cells well and faced challenges when integrating the gamma cells.

Overall, CAMEX has demonstrated competitive and robust performance across various multi-species scenarios. It can effectively handle substantial differences and multi-batch data and identify and preserve rare cell populations. Nevertheless, other baseline methods exhibit higher variability in performance, resulting in less stable outcomes.

### CAMEX uncovers the conserved differentiation process in the testis across 11 species

The testis is a unique organ with the highest number of tissue-specific genes in its transcriptome, and sperm development represents a continuous and conserved process [41]. Consequently, comparing these continuous and critical biological processes across different species is essential for evolutionary biology. Here, we collected and utilized the scRNA-seq testis data from 11 species, encompassing five distinct cell types, i.e., somatic, Sertoli, spermatogonia, spermatocytes, and spermatids [42]. Although this dataset originates from the same source and exhibits consistent cell types (**Supplementary Fig. S4a)**, integrating it across 11 species remains a significant challenge due to the scarcity of one-to-one homologous genes shared among them (**Supplementary Fig. S4b)**. Worth noting is that the original research only merges scRNA-seq data from seven primate datasets via LIGER (v.0.5.0) [24] based on primate one-to-one orthologues from Ensembl release 87 [42].

Firstly, we integrated the testis scRNA-seq data across 11 species using CAMEX and other baseline methods. In this scenario, only CAMEX successfully preserved the continuous trajectory of sperm differentiation and separated distinct clusters corresponding to different cell types (**Fig. 3a, b**). Analyzing the results of CAMEX, we observed that the Platypus dataset almost entirely lacked Sertoli cells, consistent with the original findings (**Supplementary Fig. S5a, b)**. PyLiger partially retained the developmental trajectory but failed to integrate spermatids and spermatocytes, erroneously merging somatic cells with spermatids. scVI struggled to effectively integrate sertoli and spermatocyte cells from chickens with those from other species. Harmony failed to integrate Sertoli cells from opossums and spermatocytes from chickens. Scanorama achieved the worst integration results.

**Figure 3.**
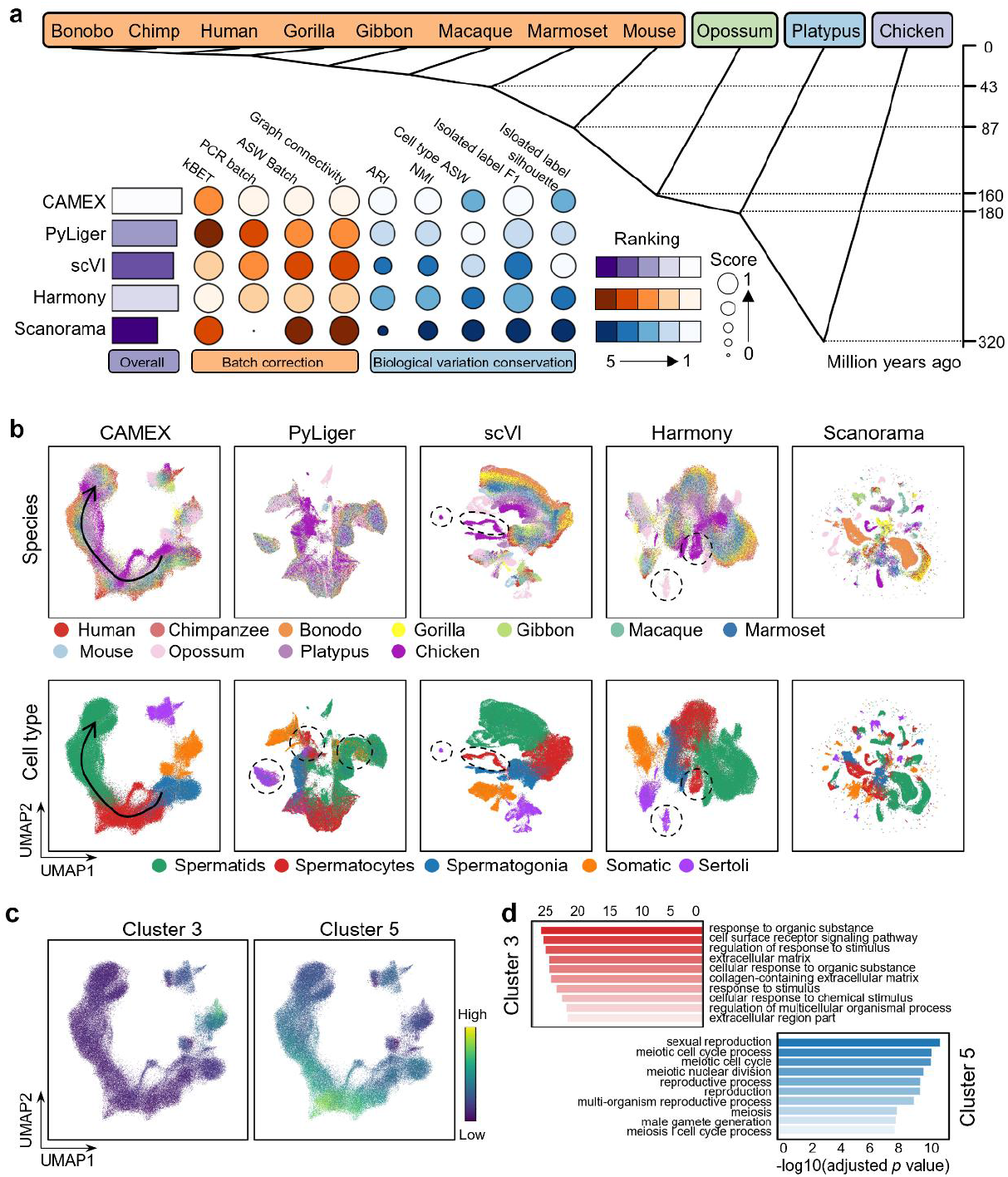
CAMEX achieved superior integration performance and preserved the differentiation process in the testis dataset across 11 species. **a**, Overall score evaluated on testis and phylogenetic tree generated by TimeTree (v.5) (http://www.timetree.org/). **b**, UMAP visualization of a joint embedding space on the testis dataset. Colors denote species annotations (first row) and cell type annotations (second row). **c**, UMAP visualization of CAMEX integration result where color denotes the mean embedding of gene clusters 3 and 5 generated by CAMEX. **d**, Gene enrichment analysis of clusters 3 and 5 using the gprofiler package.

Secondly, given that the cell subtypes from seven primate species have been manually annotated, we leveraged the cell embeddings obtained from CAMEX. Using the human dataset as a reference, we applied a simple k-Nearest Neighbors (kNN) classifier to annotate the subtypes for other primate cells. The predicted results are almost consistent with the manual annotations (normalized confusion matrix) (**Supplementary Fig. S6a**). Additionally, we annotated the cell subtypes of four other non-primate species. Cell subtypes exhibit similarities with the annotated major population (**Supplementary Fig. S7a**). CAMEX identified some cells in the platypus dataset as Sertoli cells. Through further differential analysis, we identified differentially expressed genes (DEGs) (**Supplementary Fig. S7b**). These genes are highly relevant to spermatogenesis in the testis. For example, the CYP17A1 gene codes for a protein involved in testosterone synthesis and plays a vital role in cholesterol transport [43]. Similarly, the STAR gene encodes a protein critical for cholesterol transport across mitochondrial membranes, thereby regulating testosterone synthesis [44]. We speculate that the labeled cells may represent a rare population of potential platypus-specific Sertoli cells.

Thirdly, we obtained the mean embedding for each cell type across all species and conducted hierarchical clustering (**Supplementary Fig. S8a**). The clustering outcomes indirectly confirm the accuracy of cell integration. Notably, opossum and mouse spermatogonia were mistakenly grouped with spermatocytes. We attribute this misclassification to the relatively young age of samples in the original dataset [42]. Interestingly, spermatogonia appears more similar to somatic cells than spermatocytes and spermatids. This finding suggests that the transcriptional differences during spermatogenesis are crucial [42, 45]. At the cell type level, we established a species expression hierarchy through clustering, although it remains incomplete. This exploration lays the groundwork for further research in this direction.

Finally, we acquired cell and gene embeddings via CAMEX. By clustering the gene embeddings, we identified specific functional modules within clusters 3 and 5 (**Fig. 3c, d**). Genes in cluster 3 are predominantly expressed in Sertoli cells, and enrichment analysis revealed associations with the cell surface receptor signaling pathway and extracellular matrix. Sertoli cells play a crucial role in spermatogenesis: they facilitate the commitment of fetal germ cells to the male pathway and are vital for germ cell development [46]. Conversely, genes in cluster 5 are primarily expressed in spermatogonia, and enrichment analysis linked them to sexual reproduction, meiotic cell cycle process, meiotic cell cycle, meiotic nuclear division, and reproductive process. Meiosis is a well-known process occurring in spermatocytes and is consistent with gene function obtained by CAMEX [47]. In summary, CAMEX can integrate conserved developmental processes across multiple and distant species and provide gene embeddings that shed light on cell functions.

### CAMEX aligns various development stages of seven organs across seven different species

The evolution of gene expression during various species’ organ development remains largely unknown, and different species exhibit distinct patterns. We applied CAMEX and competing methods to integrate and align the RNA-seq data from seven organs across seven species at various developmental stages [48]. Remarkably, only CAMEX successfully preserved the continuous development trajectory and separated distinct clusters corresponding to different organs (**Fig. 4a, Supplementary Figs. S9a, S10a**). We noted that the cerebellum and brain were well integrated due to their abundance of neurons and glial cells [49], while the testis and ovary, being gonadal cells in early development [50], exhibited higher similarity. Although Harmony effectively separated different organs while mixing data from various species, it failed to maintain continuous developmental trajectories (**Supplementary Fig. S9b**). scVI and Scanorama demonstrated subpar performance.

**Figure 4.**
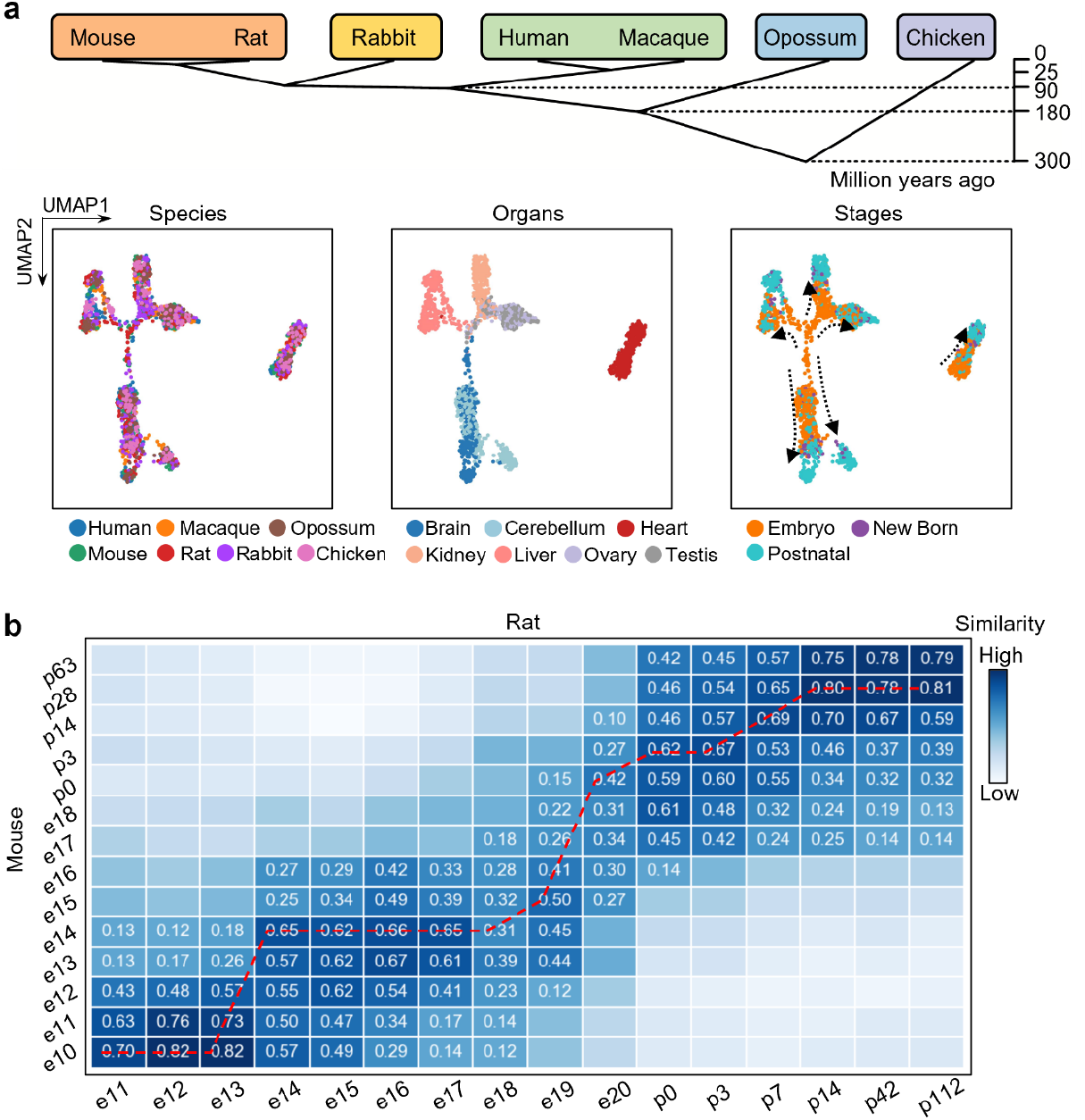
CAMEX aligns various development stages of seven organs across seven different species. **a**, Phylogenetic tree created using TimeTree (v.5) (http://www.timetree.org/). UMAP visualization of joint embedding space on this multi-species dataset, encompassing various organs and developmental stages. Cells are colored by the species (left), organs (middle), and stages (right). **b**, Similarities between all organs in mice and rats with the mouse as a reference, ensuring their developmental times were well aligned.

Most model organisms exhibit unique developmental timelines mapped to the Carnegie stages from gamy to birth. After obtaining organ embeddings from the integration results of CAMEX, we calculated the similarity of these organ embeddings, allowing us to establish correspondence across diverse species. Initially, we selected rats and mice as our subjects and sampled from embryonic day 11 (e11) to postnatal day 112 (p112) for rats and from e10 to p63 for mice. CAMEX aligned the developmental stages of mice and rats within the Carnegie stages, consistent with existing literature, and even predicted their stages beyond the specified ranges (**Fig. 4b, Supplementary Table 2**). Additionally, we employed CAMEX to analyze the brain transcriptome data from mice and opossums at multiple developmental stages. While the mouse brain data spanned from e10 to p63, the opossum brain data covered e13.5 to p180. Notably, the opossum central nervous system, including the brain, is markedly premature at birth (p0), resembling an embryonic e11 mouse, as confirmed by the similarity of development time calculated by CAMEX (**Supplementary Fig. S10b**) and previous study [51].

### CAMEX could achieve more accurate integration and annotation performance in both relatives and distant species

Integrating and annotating distant species presents a challenging task due to the tendency to share fewer orthologous genes, resulting in more specific cell-type markers. Here, we collected four scRNA-seq data from four species, i.e., adult human visual cortex, frontal cortex, cerebellum [49], mouse neocortex [52], and lizard and turtle pallium [53], which were profiled from three technology platforms. Each dataset has a specific cell type that is absent in the datasets of other species (**Supplementary Fig. S11a-c**). CAMEX, Harmony, scVI, and PyLiger achieved favorable integration performance, with CAMEX receiving the highest overall score (**Fig. 5a, Supplementary Fig. S11d**). We noted there existed instances of over-integration or under-integration in both Harmony and scVI results. For example, in the Harmony results, human astrocytes and microglial cells failed to integrate with lizard and turtle. Neural progenitor cells from turtles and lizards were over-integrated with excitatory neurons. In the scVI results, microglial cells from humans, mice, lizards, and turtles are not mixed up correctly. Similarly, PyLiger failed to integrate astrocytes properly, and a subset of astrocytes, oligodendrocyte precursors, and neural progenitors were erroneously integrated into the excitatory neuron cluster.

**Figure 5.**
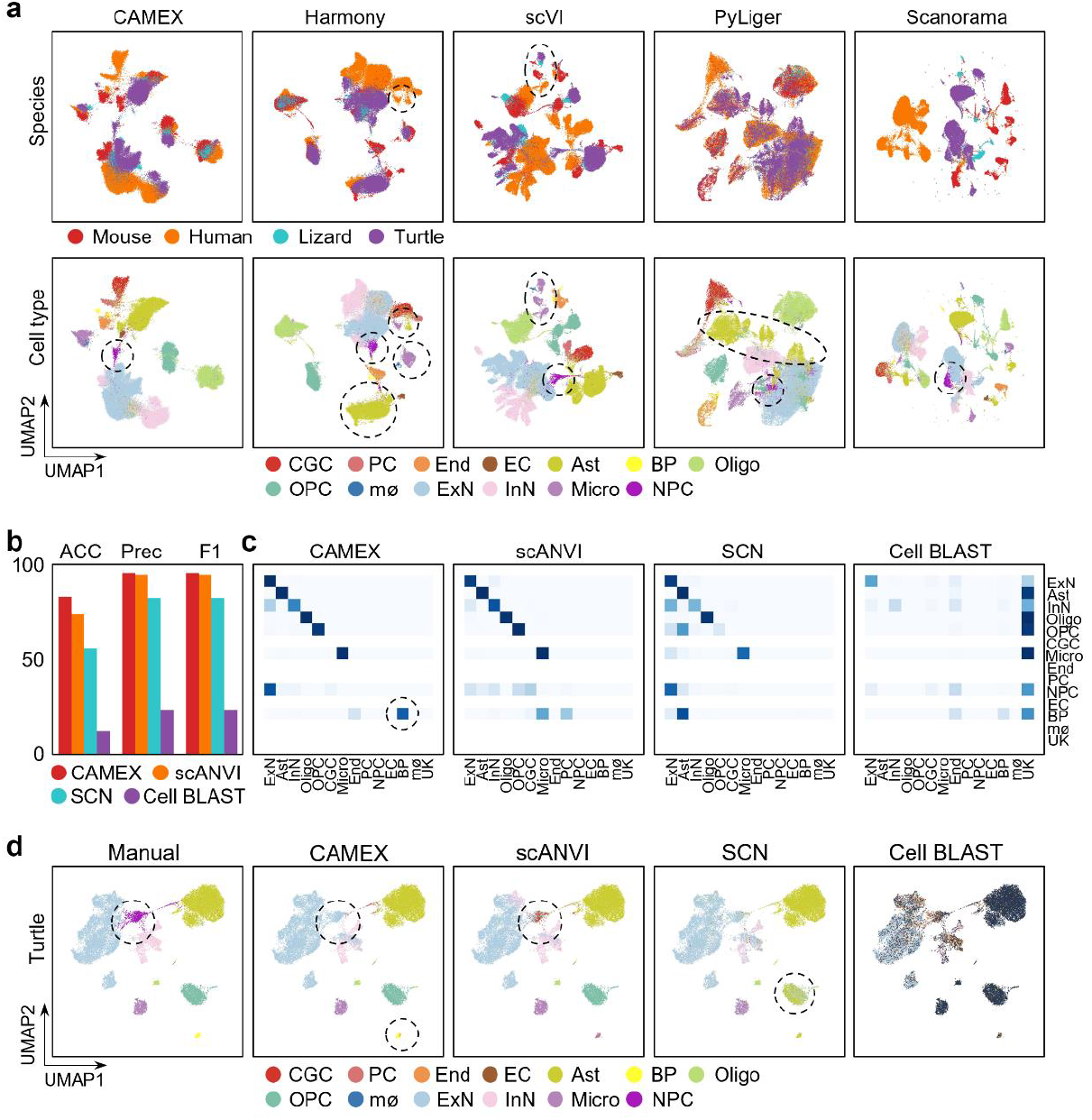
CAMEX integrates and annotates relatives and distant species cortex scRNA-seq dataset. **a**, UMAP visualization of joint embedding space. Colors denote species annotations (first row) and cell type annotations (second row). The cell types are abbreviated as follows: Astrocyte (Ast), Brain Pericyte (BP), Endothelial (End), Excitatory Neuron (ExN), Inhibitory Neuron (InN), Microglial cell (Micro), Neural Progenitor Cell (NPC), Oligodendrocyte (Oligo), Oligodendrocyte Precursor Cell (OPC), Purkinje cell (PC), Cerebellar Granule Cell (CGC), Ependymal cell (EC), Macrophage (mø), UnKnown (UK). **b**, ACC, PRC, and F1 scores achieved by CAMEX and baseline methods on the turtle dataset. **c**, Row-normalized confusion matrix of CAMEX and baseline methods on turtle dataset. **d**, UMAP visualization of turtle dataset. Colors denote manual and predicted annotations.

Traditional cell-type annotation methods encounter challenges when attempting to cluster different cell types into a single group, especially when analyzing non-model species lacking prior knowledge of cell-type biomarkers. CAMEX alleviates this issue by leveraging many-to-many homology relationships, significantly enhancing the annotation accuracy in both relatives and distant species. In this scenario, we used the human dataset as the reference (with cell labels) and applied CAMEX to annotate the other three datasets (without cell lables), particularly those from lizard and turtle samples. CAMEX demonstrated superior performance compared to baseline methods, as evaluated by metrics including ACC, Prec, and F1 score (**Fig. 5b-c, Supplementary Fig. S12a-f**). CAMEX’s annotations were accurate for most cell types, including smaller cell subgroups like brain pericytes in turtles (**Fig. 5c-d**). CAMEX identified a cluster of unknown cell types in the query dataset as excited neurons. These cells were originally defined as neural progenitor cells in the literature but can be further categorized into other mature neuron types. Previous research has shown that excitatory stimuli directly impact adult hippocampal neural progenitors, promoting neuron production [54]. We can discern this neural progenitor cell cluster clearly through unsupervised integration results (**Fig. 5a**). In contrast, scANVI, while effective at distinguishing major cell groups, mistakenly labeled the endothelial as microglial cells in the mouse dataset. SCN also exhibited slightly weaker annotation capabilities, misidentifying endothelial as microglial cells in the mouse dataset and oligodendrocyte precursor cells as astrocytes in the turtle dataset. Due to the distant genetic relationship between humans and lizards, CellBlast achieved lower accuracy due to majority rejection and the ‘unknown’ assignment. Overall, leveraging many-to-many homologous genes, CAMEX outperforms other methods in integrating and annotating cell types across remote species.

### CAMEX facilitates the discovery of new populations and markers in the primate granular dorsolateral prefrontal cortex (DLPFC) data

The discovery of new cell populations and gene markers holds immense importance in understanding evolutionary organisms and identifying novel targets. The granular dorsolateral prefrontal cortex, unique to primates in evolution, plays a pivotal role in cognitive processes. We investigated specific cell types and gene markers within the collected DLPFC dataset across four species, i.e., humans, rhesuses, marmosets, and chimpanzees [55]. We integrated all the cells in the DLPFC dataset among the four species by CAMEX. Almost all the cell types match accurately, and the major classes (e.g., excitatory neurons, inhibitory neurons, non-neural cells, and glial cells) of cell types are relatively conserved, indicating the capability of CAMEX for cross-species integration (**Fig. 6a, Supplementary Fig. S13a**).

**Figure 6.**
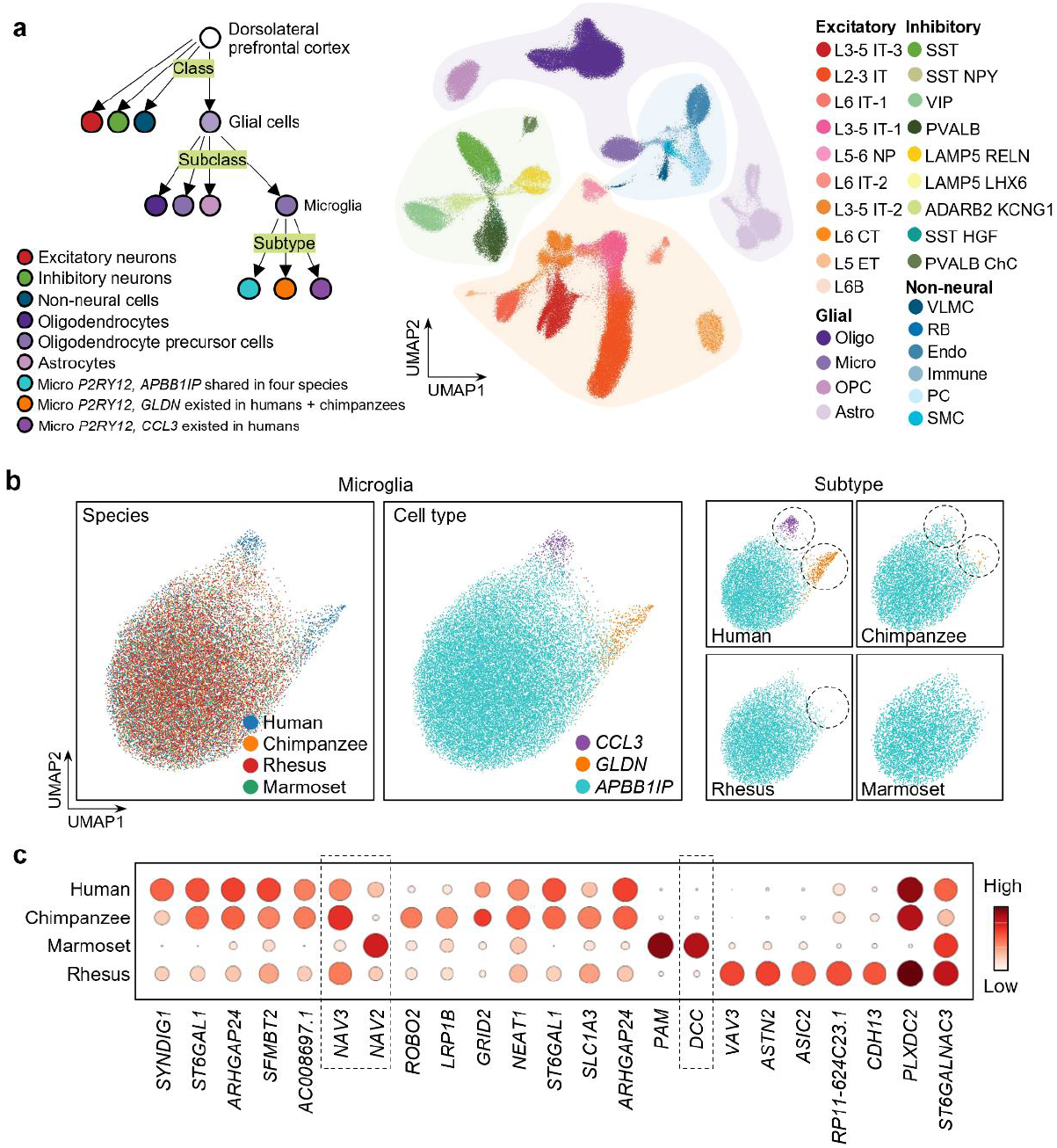
Integrating the dorsolateral prefrontal cortex (DLPFC) dataset of four species, i.e., chimpanzees, marmosets, rhesus, and humans by CAMEX. **a**, The class and subclass of cell types in DLPFC of four species, i.e., human, chimpanzee, marmoset, and rhesus. Note that two kinds of subtypes in the microglial are species specifically. UMAP visualization of a joint embedding space on the DLPFC dataset across species. Colors denote cell type annotations. **b**, UMAP visualization of a joint embedding space on microglial. **c**, Dot plots showing the differential expression genes in different species in the APBB1IP subtype.

We simulated the following scenario, assuming we were familiar with one species or model organism. We could use its comprehensive reference scRNA-seq atlas to transfer the annotations to other species and reveal some species-specific cell populations for deciphering species divergence. Considering that during the process of nervous system evolution from a diffuse to a centralized form, glial cells and neurons co-evolved, glial cells, in particular, diversified into a more varied family of cells than the latter [56]. Here, we use humans as a reference (training set). Glial cells have four subclasses, i.e., astrocyte, oligodendrocyte, oligodendrocyte precursor cell (OPC), and microglia cells. Early studies indicated that OPC and oligodendrocytes among four species shared all subtypes, and the annotation embedding also confirmed that (**Supplementary Fig. S13b**). As to astrocytes, CAMEX did not isolate the *AQP4 OSMR* subtype that existed in humans and chimpanzees (**Supplementary Fig. S13b**). This might be due to the low proportion of this subtype in the training set (<0.25%). In addition, microglia are more diverse, and those microglial cells are divided into three distinct cell subtypes, i.e., *APBB1IP, CCL3*, and *GLDN* clusters. Specifically, *CCL3* is exclusive to humans, *GLDN* is found in humans and chimpanzees, and *APBB1IP* appeared across four species (**Fig. 6a left**). Leveraging CAMEX, we mapped the other three species query into the human reference dataset, resulting in the distinct isolation of two subtypes (**Fig. 6b, Supplementary Fig. S14a, b**). We discovered that *CCL3* and *GLDN* subtypes were present in chimpanzees and rhesuses, which is inconsistent with previous claims. In humans, chimpanzees, macaques, and marmosets, the proportion of these two microglia subtypes declined sequentially, which aligns with the evolutionary distances between species. This intriguing phenomenon suggests a positive correlation between the proportion of subtypes and evolution.

Moreover, we conducted differential analysis for each subtype, comparing them with other clusters to identify subtype-specific genes (**Supplementary Fig. S15a, b**). The differential genes associated with the *CCL3* subtype primarily function in immunity. For instance, *CD83* encodes a protein known as CD83, which stabilizes MHC II, costimulatory molecules, and CD28 in the cell membrane by counteracting E3 ubiquitin ligases [57, 58]. On the other hand, the differential genes within the *GLDN* subtype are primarily involved in signal transduction. SPP1 (secreted phosphoprotein 1) is an extracellular protein closely linked to tumor biology, including processes like proliferation, migration, and invasion [59, 60].

In addition, we analyzed species differences. Even within the *APBB1IP* subtype, we observed abundant differential genes between species (**Fig. 6c**). *NAV2* and *NAV3* belong to the neuronal navigator family and are predominantly expressed in the nervous system. They likely play roles in axon guidance, cell migration, and neurodevelopment [61]. PAM is involved in the amidation process of peptide hormones and neurotransmitters and is crucial for regulating their activity and stability [62]. Overall, the human and chimpanzee transcriptomes exhibit a high consistency at the single-cell resolution and can serve as model organisms for exploring disease mechanisms or drug development. However, focusing on more species-specific genes is essential when studying marmoset and rhesus.

## Discussion

The scRNA-seq data is becoming increasingly popular, encompassing a wider range of species. Comprehensive cross-species comparisons and analyses at the single-cell resolution can significantly enhance our understanding of cellular origins and evolutionary mechanisms. By exploring conserved and specific features of cell states, one can gain insights into fundamental evolutionary processes and how cellular diversity correlates with physiological connections across species. Consequently, there is a pressing demand for robust tools to integrate multi-species datasets.

While previous approaches have made significant efforts toward cross-species integration and annotation, their widespread applicability to current multi-species data remains limited due to various constraints. For instance, integration algorithms designed for same-species data often focus solely on one-to-one homologous genes when applied to multi-species datasets. Unfortunately, these algorithms falter when dealing with remote species due to the distinct number of gene copies involved in evolutionary processes [63]. Another type of algorithm constructs correlations between diverse species based on gene sets (pathway and module) [20]. However, the availability of prior knowledge regarding gene sets is often restricted to model organisms, rendering this algorithm ineffective when applied to non-model organisms.

Here, we introduce CAMEX for integrating, aligning, and annotating multi-species scRNA-seq data using heterogeneous graph neural networks. We treat the dataset for each species as a bipartite graph and connect these bipartite graphs through many-to-many homology relationships. This design allows us to leverage homologous relationships as much as possible. For integration tasks, CAMEX employs an unsupervised encoder-decoder architecture. For annotation tasks, it utilizes a semi-supervised encoder-classifier architecture. We demonstrate the effectiveness of CAMEX on multiple single-cell datasets from different organs across various species.

Additionally, we integrate sperm development data from up to 11 species and validate somatic and spermatocyte functions through gene embedding. We align multi-species data across different developmental stages, which extends beyond the Carnegie Stage. The cell embeddings obtained by CAMEX facilitate the discovery of species-specific cell subtypes and novel marker genes. CAMEX also serves as a robust cell annotation tool, showcasing competitive performance compared to baseline methods across multiple datasets. In short, we expect CAMEX to become a significant tool for understanding the conservation and diversification of cell types across species, revealing fundamental evolutionary processes at the single-cell transcriptomic resolution.

While CAMEX demonstrates significant utility, it also exhibits some limitations that warrant improvement. First, the similarity of homologous genes can be obtained through sequence alignment. However, other methods for calculating similarity have not been considered in CAMEX. Fine-tuning this aspect could enhance its performance. Second, despite incorporating many-to-many homologous relationships into the graph structure, CAMEX still relies on one-to-one homologous genes during initializing cell node features. This limitation impacts its implementation and performance when integrating data across highly diverse species. Third, CAMEX leverages cell and gene embeddings for interpretability. However, the interpretability analysis based on graph neural networks remains an ongoing challenge. Exploring computational methods to elucidate biological phenomena, such as inferring gene regulation networks and considering subgraph structures as potential pathways, could yield unexpected insights. Last, as technology evolves, we envision CAMEX broadening its scope to encompass multi-omics and spatial transcriptomics. This approach would provide a more comprehensive map for species origin and evolution by leveraging multi-dimensional information.

## Methods CAMEX model

### Heterogeneous graph of cells and genes

Each gene expression matrix with *N* cells and *M* genes can be defined as *X* = (*x*_1_, *x*_2_, …, *x*_*N*_)^*T*^ ∈ *R*^*N*×*M*^. After normalization, log-transformation, selection of highly variable genes, clustering, and obtaining differential genes, we convert the expression profile into a cell-to-gene bipartite diagram *A*, where cell *n* links gene *m* if and only if cell *n* expresses gene *m*. The cell *n*_1_ and cell *n*_2_ will be connected if they are neighbors calculated by the kNN algorithm in the PCA space. In addition, we added a self-loop to each cell and gene, respectively. Finally, we connected genes from different species through the many-to-many-homologous gene relationships. In short, there are two types of nodes including cells and genes, and six types of edges (relations) including ‘a cell expresses a gene’, ‘a gene is expressed by a cell’, ‘two cells that are kNN neighbors’, ‘self-loop of a cell’, ‘self-loop of a gene’ and ‘two genes that are orthologous ones’ in the heterogeneous graph.

### Heterogeneous graph neural network encoder

CAMEX adopts a heterogeneous graph neural network as the encoder inspired by the relational graph convolutional network [64]. We design six independent parameter matrices for each of the above relationships as follows. *W*_*cg*_ (a cell expresses a gene), *W*_*gc*_ (a gene is expressed by a cell), *W*_*cc*_ (two cells that are kNN neighbors), *W*_*gg*_ (two genes that are orthologous ones), *W*_*c*_ (self-loop of a cell), and *W*_*g*_ (self-loop of a gene). The encoder transforms the features of the neighbors in a non-linear way and converges to the center node according to the message-passing paradigm. The (*l* + 1)*-*th layer representation of cell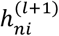 and gene 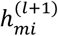 can be defined as follows:

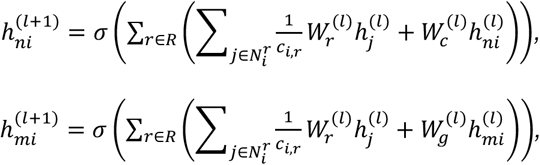

where 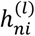 and 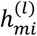 represent the *l-*th layer representation of cells and genes, respectively, 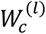 and 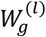 represent the *l-*th layer weight matrices of ‘self-loop of a cell’ and ‘self-loop of a gene’, respectively, *N*^*r*^ denotes the set of neighbor indices of the node (cell or gene) *i* under relations 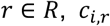 is a normalization factors, 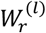 is the weight matrix of the *l-*th layer transformation of the relationship 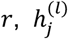 is the *l-*th layer representation of neighbor *j*, and *σ* is the activation function (i.e., Leaky ReLU here).

To reduce the number of parameters to prevent overfitting, we share weight matrices of the same relationship in different layers such as 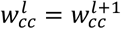.After the message passing through several layers of neural networks, the cell and gene representation can be mapped into the same embedding space.

### Edge and feature reconstruction decoder

When integrating different scRNA-seq data in an unsupervised manner, we employ two decoders to reconstruct edges and features, respectively. Specifically, the dot-product decoder aims to reconstruct the edges of the input heterogeneous graph. Edge predictions in the heterogeneous graph are generated using the dot product of cell and gene embeddings:

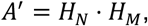

where *H*_*N*_ and *H*_*M*_ are the cell and gene embeddings, respectively, and *A*^′^ is the reconstructed adjacency matrix.

We measure the quality of each edge in the adjacency matrix using cross-entropy loss:

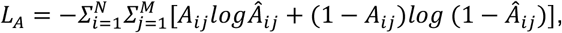

where *A*_*ij*_ is the edge between the *i-*th cell and the *j-*th gene in the original graph.

The feature reconstruction decoder aims to reconstruct cell features through a fully connected layer:

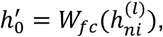

where 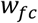 is the weight of a fully connected neural network and *h* is the reconstructed cell feature.

We utilize the mean squared error to measure the distance between the original *h*_0_ and the predicted 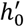:

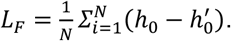

### Graph attention classifier

Cell labels are not mandatory during the integration task. However, during annotation, it is essential to have one or more reference datasets with pre-defined manual annotations. Consequently, the model could be trained in a semi-supervised manner. The cell annotations provided by the reference dataset can be denoted as: *Y* = (*y*_1_, *y*_2_, …, *y*_*Nr*_) ∈ *R*^*c*^, where *Nr* represents the number of cells in the reference dataset and *C* denotes the set of cell types. When performing cell annotation tasks, cell embeddings can be input into a GAT classifier. Specifically, the attention coefficients are first computed as follows:

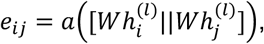

where 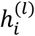 and 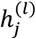 is the *l-*th layer representations of cells *i* and *j*. Shared parameter *W* maps cell embedding to a common space. The concatenation operation (denoted by ||) is applied. Finally, the function *a* maps the concatenated result to a scalar representing the relevance between nodes *i* and *j*. Subsequently, the relevance scores are normalized to obtain attention coefficients:

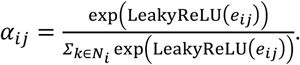

We use the multi-head self-attention mechanism to predict the final results:

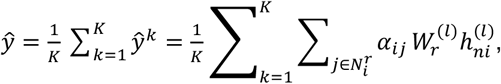

where *K* is the total number of attention-heads, set as eight by default. We optimize the semi-supervised model using the cross-entropy loss function.

### Training details. (1) Graph sampling

For large-scale data with a large number of cells or involving more species, we partition the heterogeneous graph into multiple subgraphs and train the graph neural network using a mini-batch approach. We implement this using the Deep Graph Library (DGL) [65].

**(2) Checkpoint selection:** In deep learning model training, checkpoints are commonly employed to prevent overfitting. However, in the context of unsupervised integration and semi-supervised annotation tasks, these checkpoints can sometimes lead to over-correction or over-fitting. To address this, we introduce a novel metric for evaluating checkpoints. Specifically, we cluster cells based on their embeddings and assess whether to halt training or revert to a checkpoint using the clustering metric ARI.

### Baseline methods

#### Integration methods

##### Harmony

Harmony [19] has demonstrated superior performance in various benchmarking studies for scRNA-seq data integration. We applied the Python package harmony (v0.0.9) in this study.

##### scVI

scVI [66] is an extensible Python toolkit with diverse applications. We applied the Python package scvi-tools (v0.14.6) in this study.

##### pyLiger

LIGER [24] is a robust online integration model. It has recently been re-implemented as PyLiger [23] in Python with improved computational efficiency. We applied the Python package PyLiger (v0.1.3) in this study.

##### Scanorama

Scanorama [25] is a sensitive algorithm that identifies and merges shared cell types between all datasets and accurately integrates heterogeneous scRNA-seq datasets. We applied the Python package Scanorama (v1.7.3) in this study.

#### Annotation methods

##### scANVI

scANVI [29] is a probabilistic approach to annotate cells in a semi*-* supervised way. We applied the Python package scvi-tools (v0.14.6) in this study.

##### CellBlast

CellBlast [67] is an easy-to-use Python package for cell-type annotation. We applied the Python package CellBlast (v0.5.0) in this study.

##### SingleCellNet

SingleCellNet [18] is a tree model-based R package that can annotate single-cell datasets including cross-species RNA-seq data, and the authors recently migrated this approach into Python (https://github.com/pcahan1/PySingleCellNet/).

#### Evaluation Metrics

##### Integration metrics

We group the batch correction metrics into two categories: 1) batch correction and 2) conservation of biological variance. Specifically, the batch correction score using metrics including kBET (k-nearest-neighbor Batch Effect Test) [26], PCR batch (Principal Component Regression of the batch covariate) [26], ASW batch (Average Silhouette Width across batch) [26], and Graph connectivity [14]. The conservation of biological variation score using metrics including ARI (Adjusted Rand Index) [27], NMI (Normalized Mutual Information) [28], Cell type ASW [14], ISO label F1 (Isolated label F1) [14], and ISO label silhouette (Isolated label silhouette) [14]. We implemented all the above integration metrics by scIB [14].

##### kBET

The kBET algorithm assesses whether the label composition of a cell’s k nearest neighborhood is consistent with the expected label distribution. This evaluation is performed by repeating the test on a random subset of cells, and the outcomes are aggregated as a rejection rate across all tested neighborhoods.

##### PCR batch

PCR batch quantifies the batch effect removal by comparing the variance contributions of the batch effects of datasets before integration (*Vc*_*before*_) and after integration (*Vc*_*after*_), respectively.

##### ASW Batch

ASW Batch calculates the silhouette width of cells with respect to batch labels.

##### Graph connectivity

Graph connectivity assesses whether the graph correctly connects cells of the same cell-type labels among batches.

##### ARI

The adjusted Rand index (ARI) is a common external evaluation metric for clustering. It assesses the effectiveness of clustering by calculating the number of sample pairs assigned to the same or different clusters in both the true labels and the clustering results.

##### NMI

NMI computes normalized mutual information between two clustering results.

##### Cell type ASW

Cell-type ASW evaluates ASW with respect to cell-type labels, where a higher score means that cells are closer to cells of the same cell type. **Isolated label F1:** Isolated label F1 is developed to measure the ability of integration methods to preserve dataset-specific cell types. We adopted the Scanpy pipeline to cluster cells and evaluated the cluster assignment of dataset-specific cell types based on the F1 score.

##### Isolated label silhouette

Isolated label silhouette measures the conservation of dataset-specific cell types. It evaluates the ASW of dataset-specific cell types.

##### Annotation metrics

To comprehensively compare the performance of different annotation methods, we employed three kinds of evaluation metrics, including accuracy, precision, and F1 score. Cell annotation is a multi-classification task. Considering the difference in the number of different types of cells in a dataset, all evaluation metrics are weighted. We implemented all the above annotation metrics by Scikit-learn [28].

##### Accuracy

The accuracy can be defined as follows:

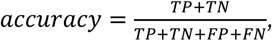

where *TP* is the number of true positives, *TN* is the number of true negatives, *FP* is the number of false positives, and *FN* is the number of false negatives.

##### Precision

The precision can be defined as follows:

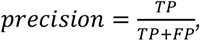

where *TP* is the number of true positives and *FP* is the number of false positives.

##### F1 score

The F1 score can be interpreted as a harmonic mean of precision and recall, where an F1 score reaches its best value at 1 and worst score at 0. The F1 can be defined as follows:

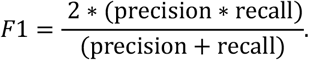

## Data Availability

The details of all datasets can be found in **Supplement Table S1**. We preprocessed all raw data following the pipeline by Scanpy [68]. For the baseline methods, we map the gene symbols to the first or reference species dataset using one-to-one homologous relationships.

## Code Availability

The source codes of CAMEX is accessible via this link on GitHub: https://github.com/zhanglabtools/CAMEX/.

## Acknowledgements

This work has been supported by the National Key Research and Development Program of China [No. 2021YFA1302500 to S.Z.], the National Natural Science Foundation of China [Nos. 32341013, 12326614 to S.Z., Nos. 62333018, 62372255,61932008, U22A2039, 62372318, 62073231 to D.S.H.], the CAS Project for Young Scientists in Basic Research [No. YSBR-034 to S.Z.], the STI 2030—Major Projects [No. 2021ZD0200403 to D.S.H.], the Key Research and Development (Digital Twin) Program of Ningbo City [Nos.2023Z226, 2023Z219 to D.S.H.], the Key Project of Science and Technology of Guangxi [No. 2021AB20147 to D.S.H.], the Guangxi Natural Science Foundation [Nos. 2022JJD170019, 2021JJA170204, 2021JJA170199 to D.S.H.] and Guangxi Science and Technology Base and Talents Special Project [2021AC19354, 2021AC19394 to D.S.H.] and the Guangxi Key Lab of Human-machine Interaction and Intelligent Decision, Guangxi Academy Sciences.

## Author Contributions

S.Z. conceived and supervised the project. Z. G. collected the datasets and developed the algorithm. Z. G., D. H. and S.Z. performed the analyses and wrote the manuscript. All authors read and approved the final manuscript.

