## Supplemental Text and Figures for "Multi-species integration, alignment and annotation of single-cell RNA-seq data with CAMEX"

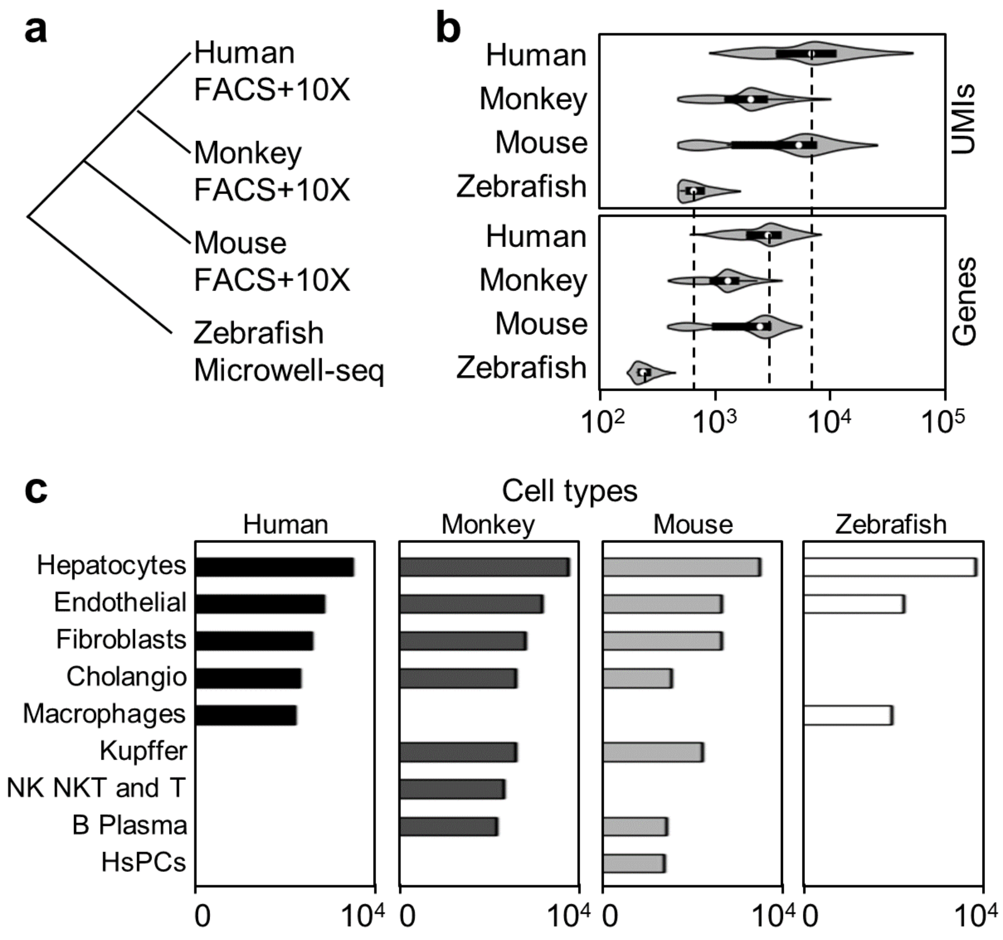


**Supplementary Figure S1.** Statistics for the liver dataset across four species. **a**, Liver scRNA-seq dataset includes samples from humans, macaques, mice, and zebrafishes. **b**, Number of detected UMIs and genes in different species. **c**, Number of cells in each cell type for each species. Each dataset comprises distinct cell types that are unique to that respective species.

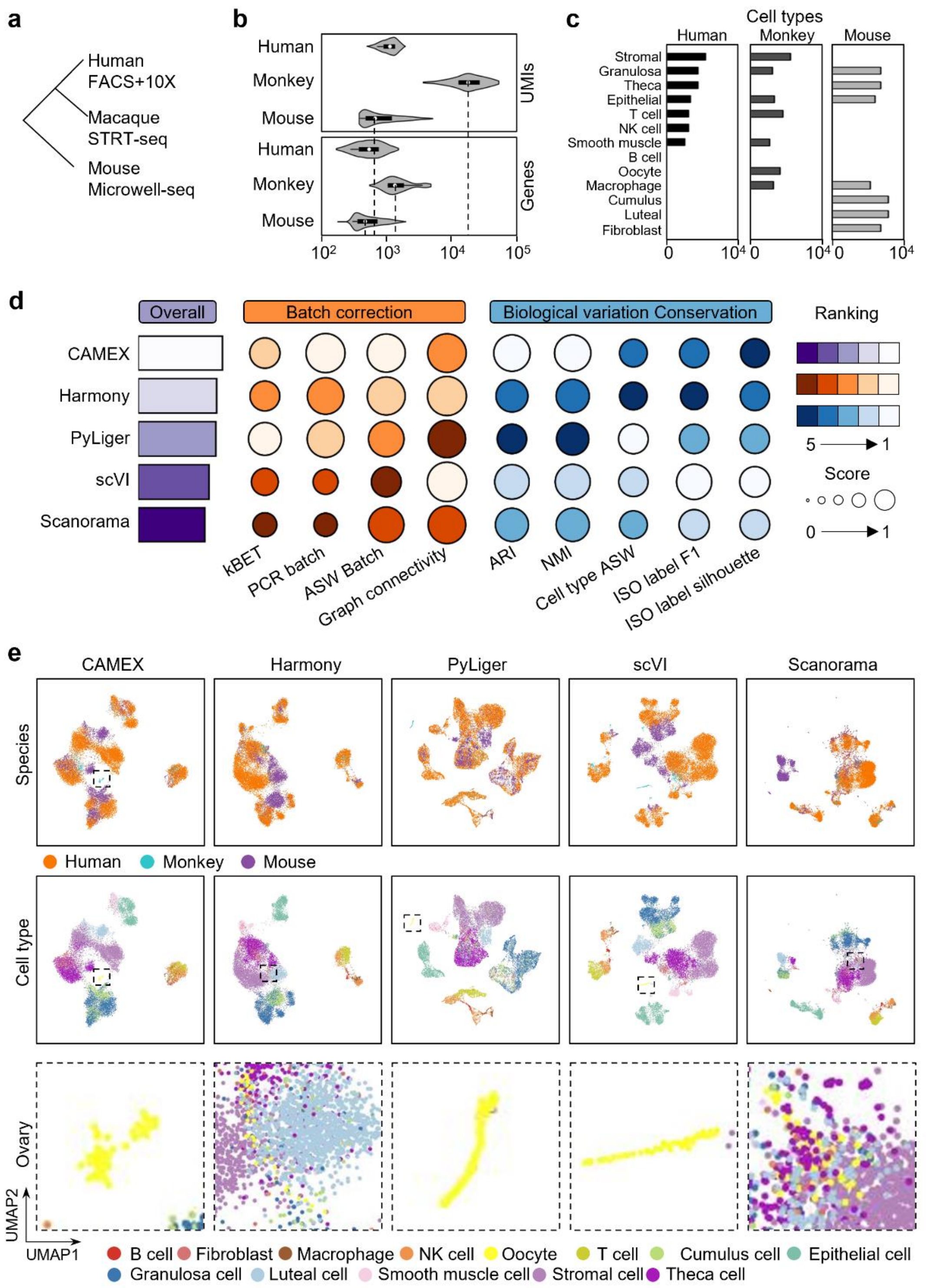


**Supplementary Figure S2.** Benchmarking CAMEX and SOTA methods on ovary dataset across three species. **a**, Ovary scRNA-seq dataset includes samples from humans, macaques, and mice. **b**, Number of detected UMIs and genes in different species. **c**, Number of cells in each cell type for each species. Each dataset comprises distinct cell types that are unique to that respective species. **d**, Overall score evaluation on the ovary. **e**, UMAP visualization of a joint embedding space on the ovary dataset. Colors represent species annotations (first row), cell type annotations (second row), and oocyte cells.

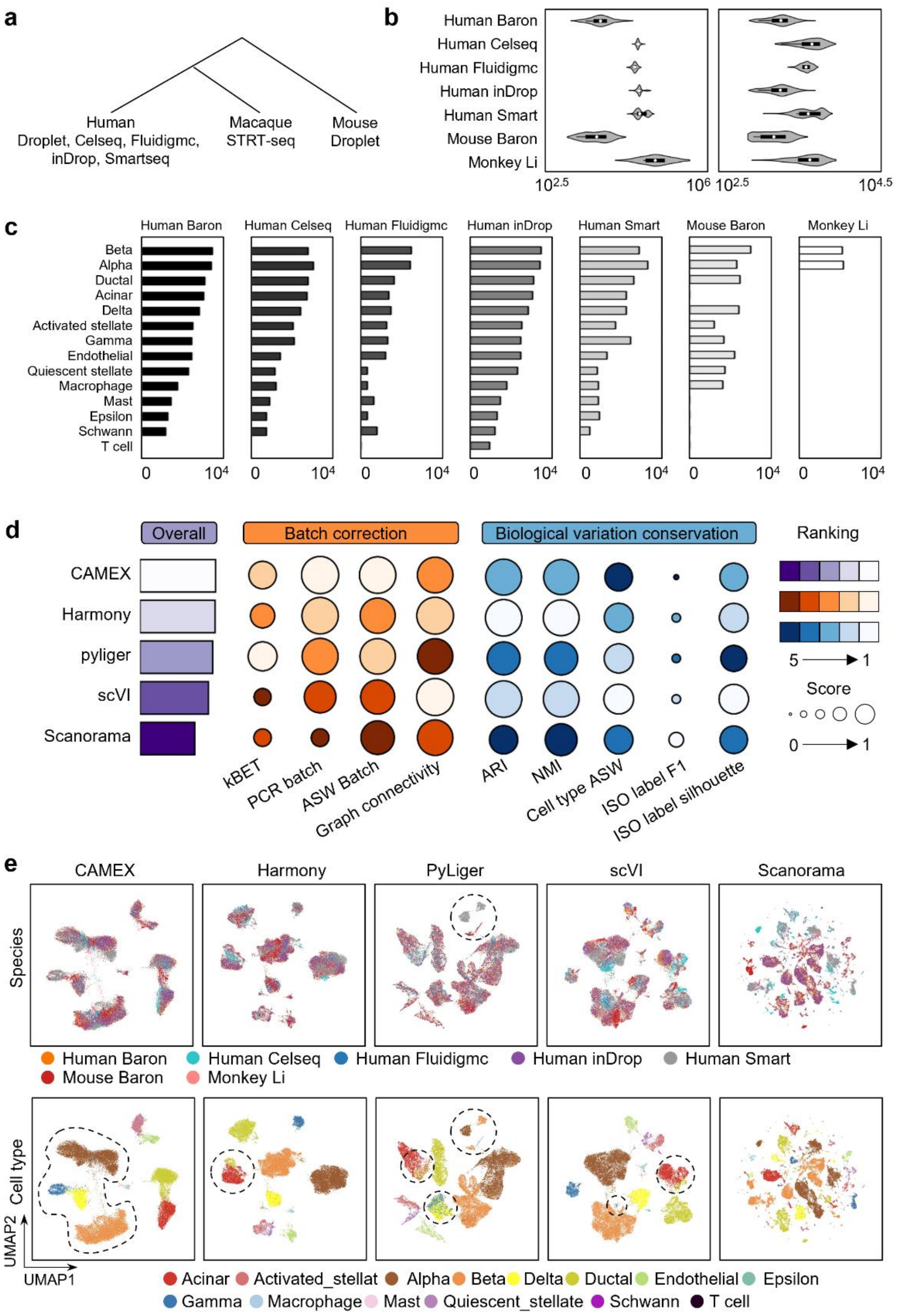


**Supplementary Figure S3.** Benchmarking CAMEX and SOTA methods on the pancreas dataset across humans, macaques, and mice. **a**, Pancreas scRNA-seq dataset includes samples from three species. **b**, Number of UMIs and genes in different species. **c**, Each dataset comprises distinct cell types that are unique to that respective species. **d**, Overall score evaluation on pancreas. **e**, UMAP visualization of a joint embedding space on the ovary dataset. Colors represent species annotations (first row), cell-type annotations (second row) and oocyte cells.

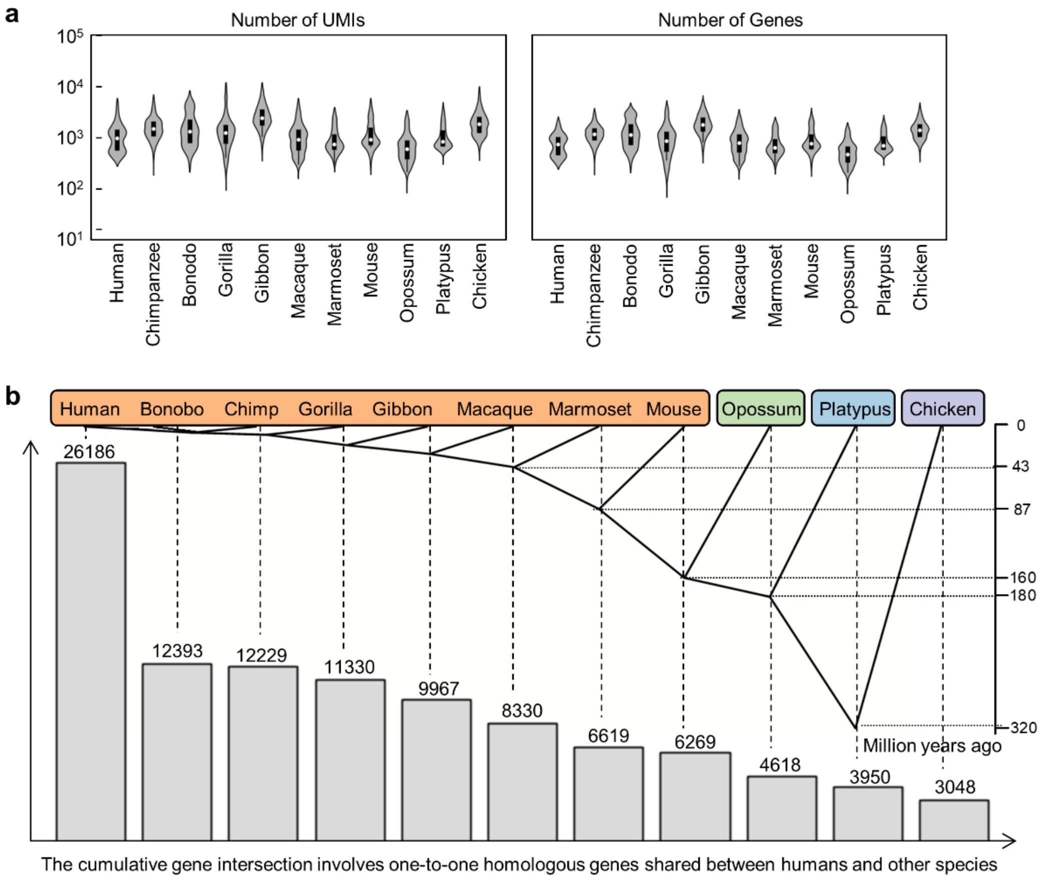


**Supplementary Figure S4. Statistics for the testis dataset across 11 species. a**, Number of the detected UMIs and genes in different species. **b**, The gene intersection involves one-to-one homologous genes shared between humans and other species. For example, the human dataset contains 26,186 genes, the number of one-to-one homologous genes between human and chimpanzee datasets is 12,393, and the number of one-to-one homologous genes among all 11 species was 3048.

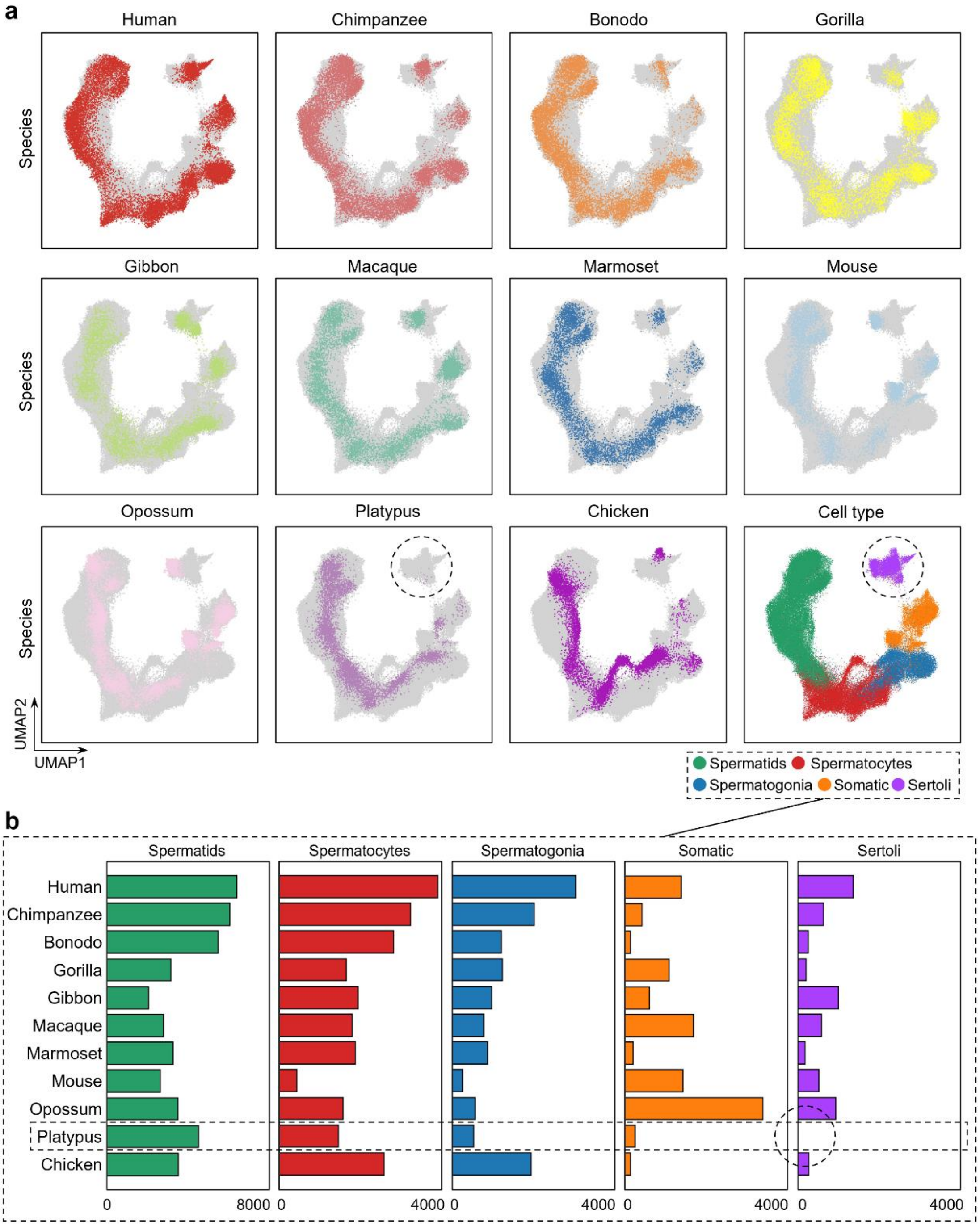


**Supplementary Figure S5.** Integration results of CAMEX on testis across 11 species. **a**, UMAP visualization of CAMEX integration result where color denote different species. **b**, Distributions of cell types across 11 species.

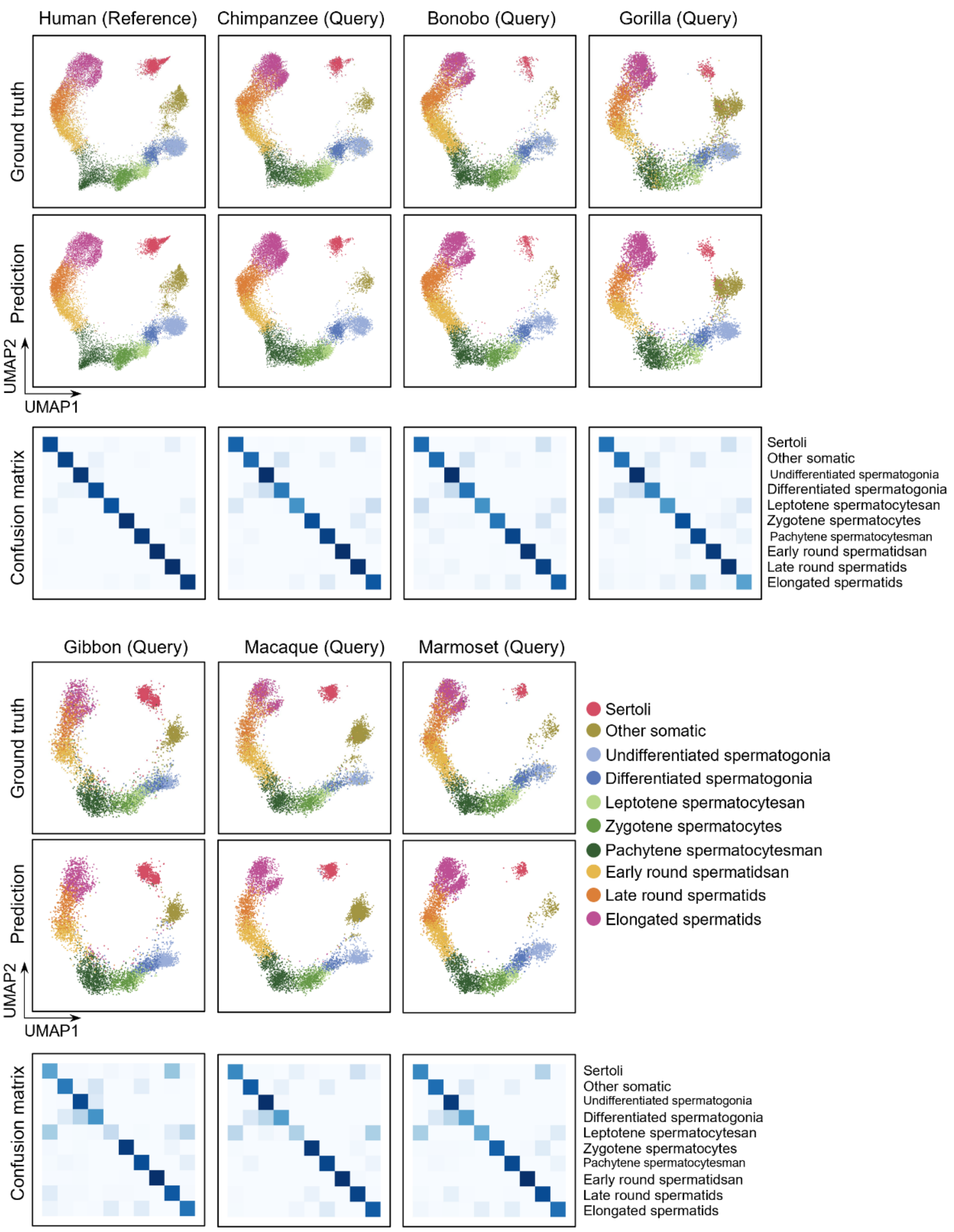


**Supplementary Figure S6.** Integration and kNN annotations of CAMEX on the 11-species testis dataset, focusing on the outcomes for seven primate species. UMAP visualization of a shared embedding space for the testis dataset. Colors correspond to manual (first row) and predicted annotations (second row) and the row-normalized confusion matrix.

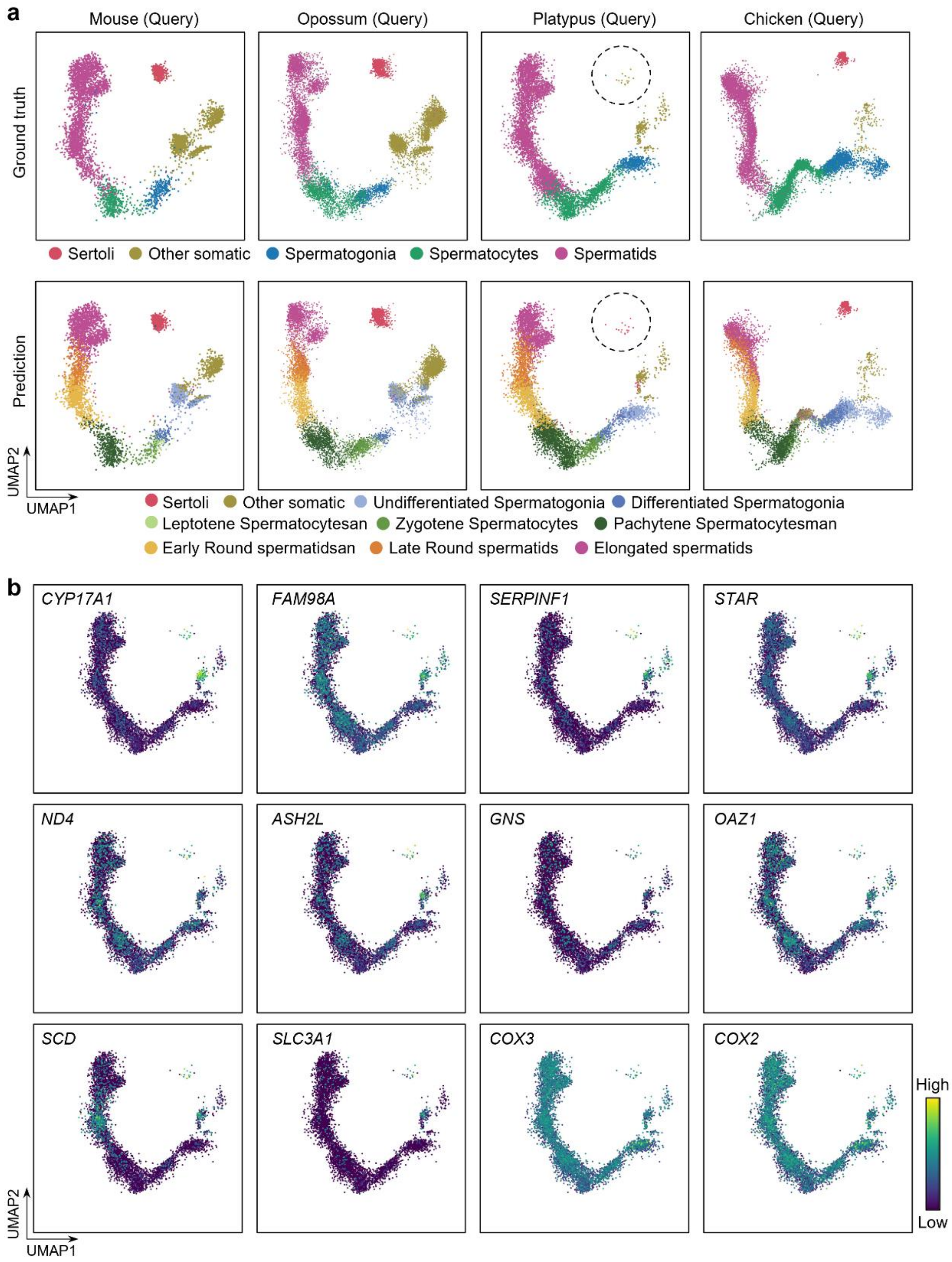


**Supplementary Figure S7.** Integration and kNN annotations CAMEX on the 11-species testis dataset, focusing on the outcomes for four non-primate species. **a**, UMAP visualization of a joint embedding space for the testis dataset. Colors correspond to manual (first row) and predicted annotations (second row). **b**, Visualizations of 12 differential genes in the platypus dataset.

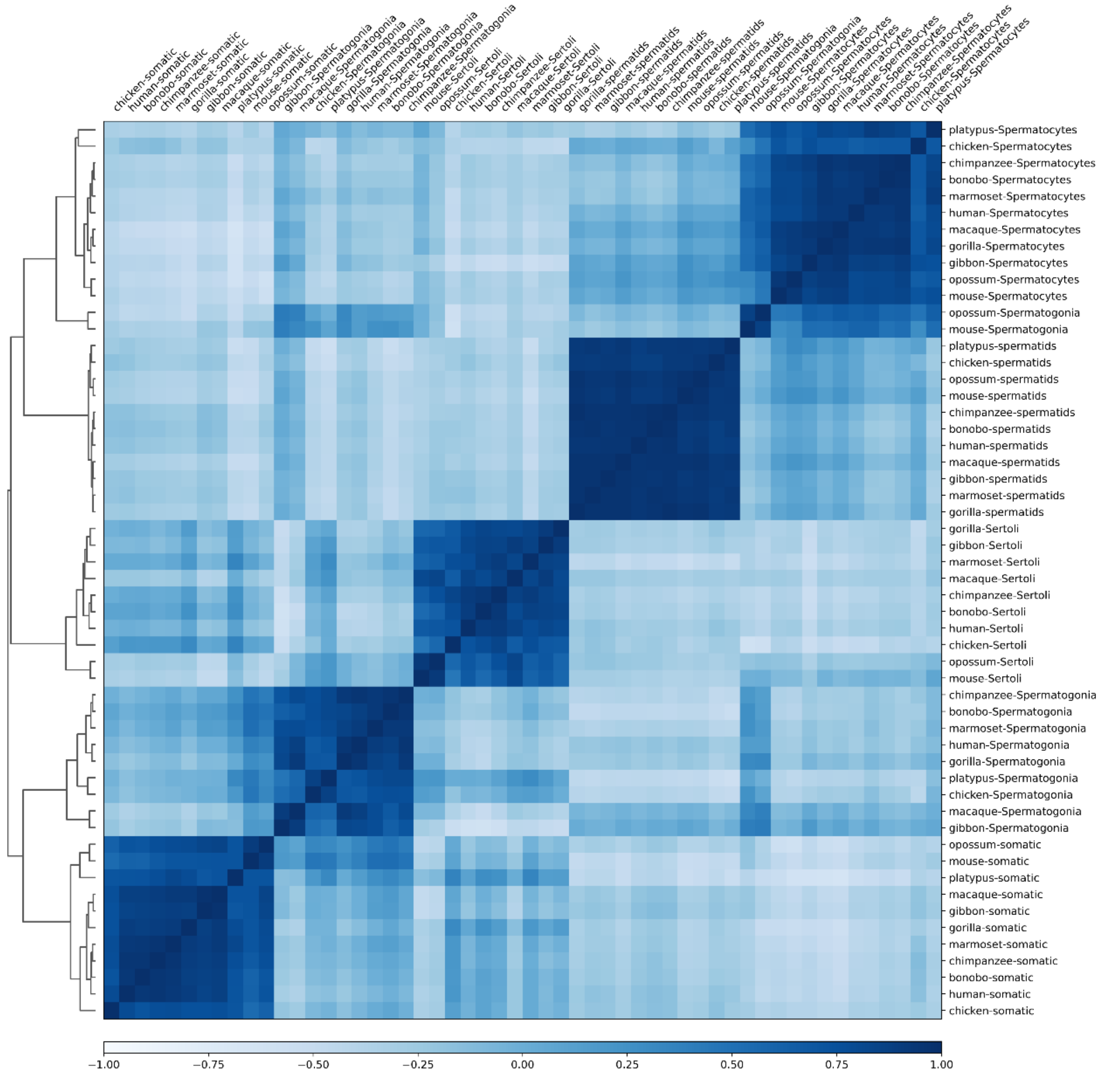


**Supplementary Figure S8.** Hierarchical clustering of the mean embedding for each cell type across all 11 species.

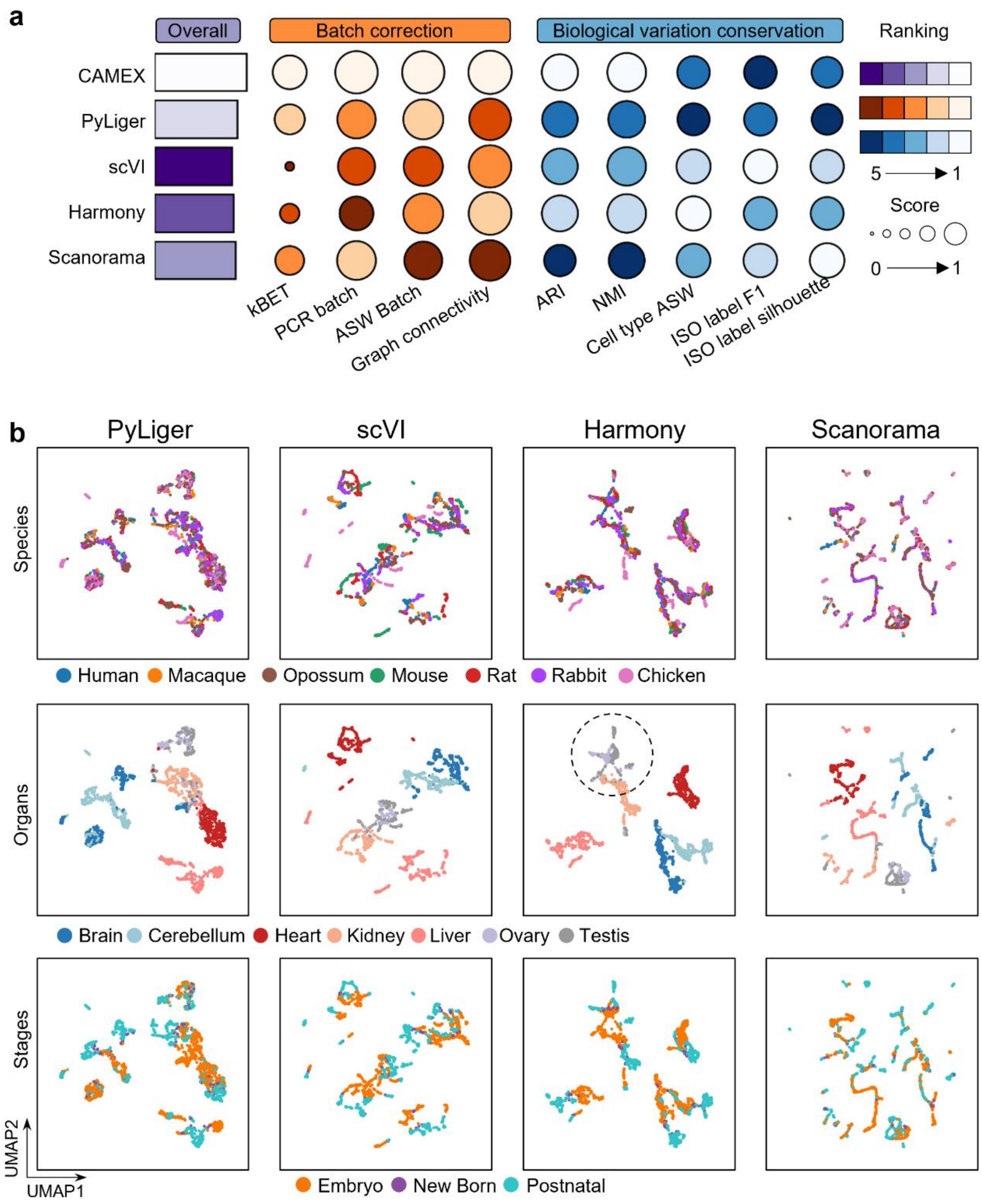


**Supplementary Figure S9.** Integration results of SOTA methods on multi-species dataset, encompassing various organs and different developmental stages. **a**, Overall score evaluation on this dataset. **b**, UMAP visualization of a joint embedding. Colors correspond to species annotations (first row), organs annotations (second row), and stages annotations (third row).

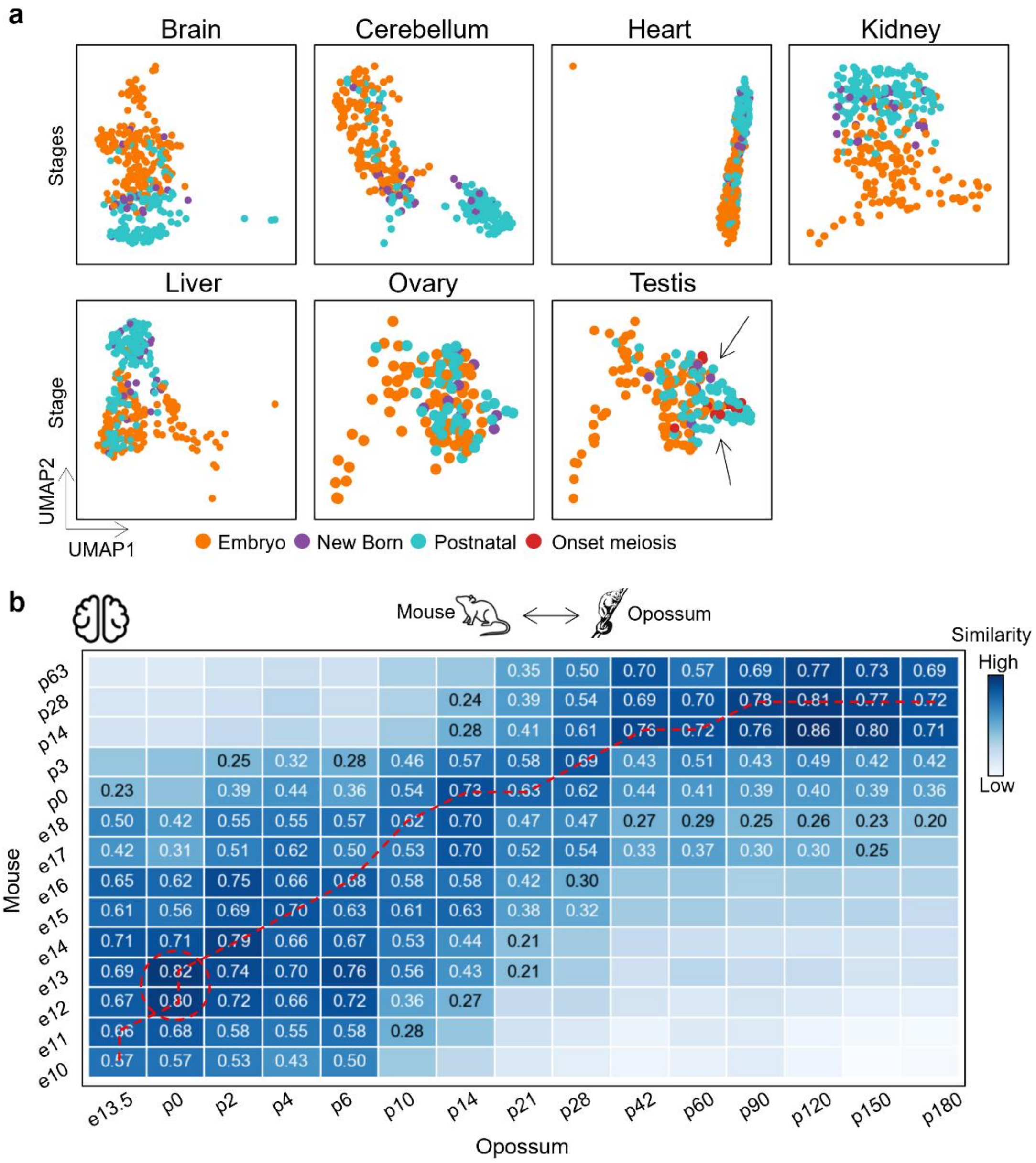


**Supplementary Figure S10.** Integration results of multi-species dataset with CAMEX, encompassing various organs and different developmental stages. **a**, UMAP visualization of a shared embedding space of each organ. Colors correspond to development stages. **b**, Similarities between brains in mice and opossums using the mouse as a reference, ensuring their developmental times were well aligned. The opossum central nervous system, including the brain, is markedly premature at birth (p0), resembling an embryonic e11 mice.

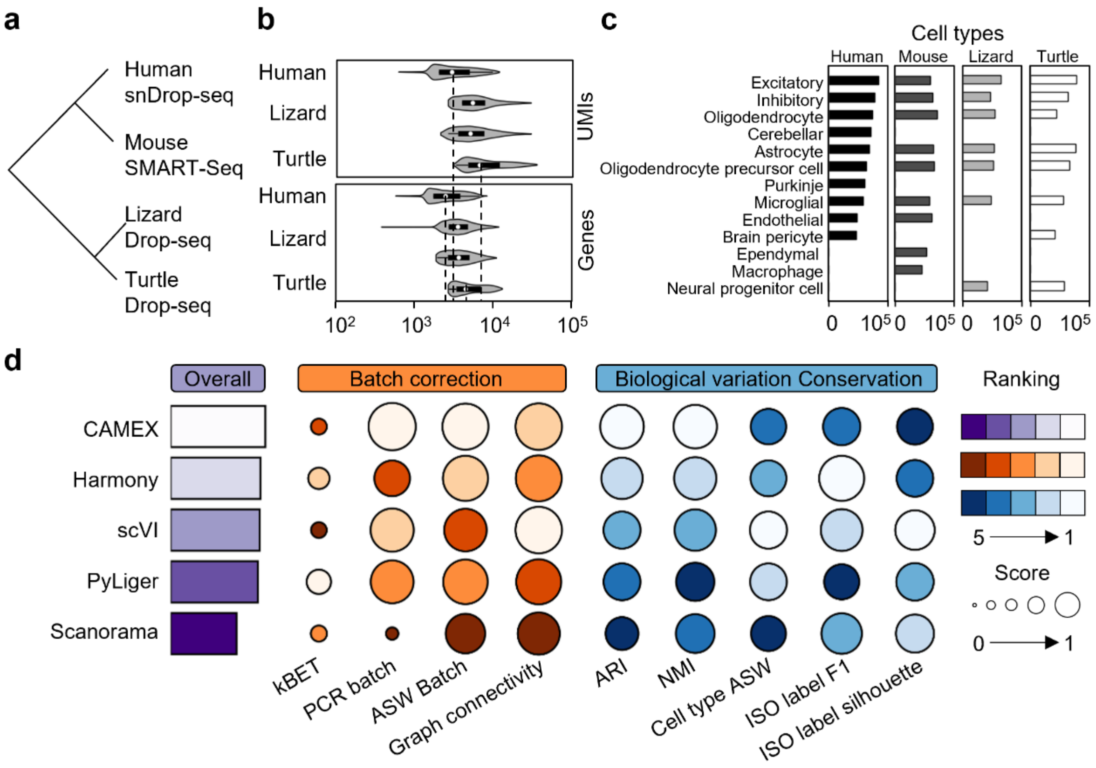


**Supplementary Figure S11.** Benchmarking CAMEX and baseline methods on the cortex dataset across four species. **a**, Cortex scRNA-seq dataset includes samples from humans, mice, lizards, and turtles. **b**, Number of detected UMIs and genes in different species. **c**, Each dataset comprises distinct cell types that are unique to that respective species. **d**, Overall score evaluation on the cortex dataset.

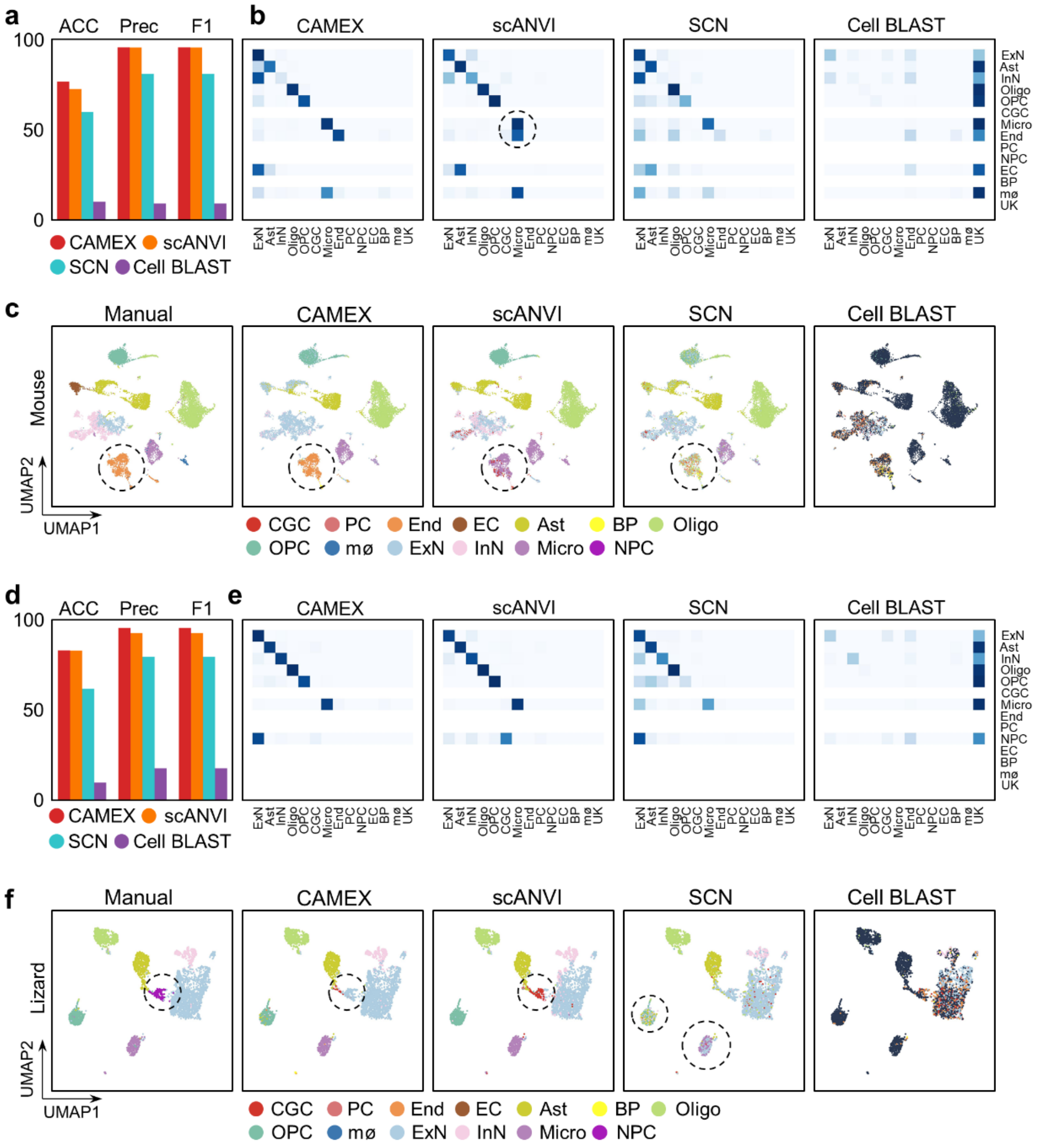


**Supplementary Figure S12.** Predicted annotation by CAMEX and baseline methods. The cell types are abbreviated as follows: Astrocyte (Ast), Brain Pericyte (BP), Endothelial (End), Excitatory Neuron (ExN), Inhibitory Neuron (InN), Microglial cell (Micro), Neural Progenitor Cell (NPC), Oligodendrocyte (Oligo), Oligodendrocyte Precursor Cell (OPC), Purkinje cell (PC), Cerebellar Granule Cell (CGC), Ependymal cell (EC), Macrophage (mø), UnKnown (UK). **a**, ACC, PRC, and F1 scores achieved by CAMEX and baseline methods on the mouse dataset. **b**, Row-normalized confusion matrix of CAMEX and baseline methods on the mouse dataset. **c**, UMAP visualization of the mouse dataset. Colors denote manual and predicted annotations. **d**, ACC, PRC, and F1 scores achieved by CAMEX and baseline methods on the lizard dataset. **e**, Row-normalized confusion matrix of CAMEX and baseline methods on the lizard dataset. **f**, UMAP visualization of the lizard dataset. Colors denote manual and predicted annotations.

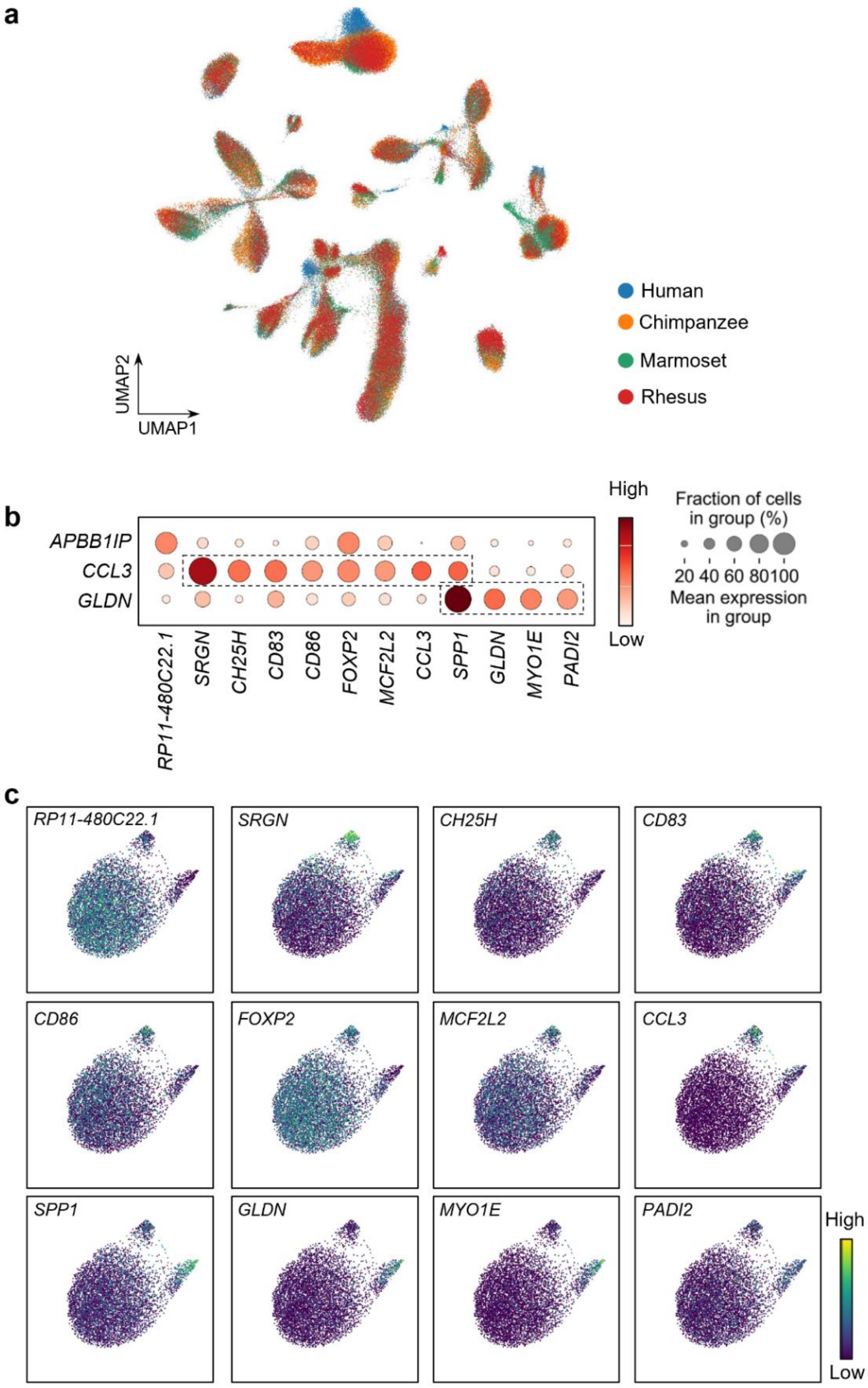


**Supplementary Figure S13.** Integrating the dorsolateral prefrontal cortex data of four species, i.e., chimpanzees, marmosets, rhesus, and humans, taking humans as a reference using CAMEX. **a**, UMAP visualization of the whole dataset across four species. **b**, Dot plots showing the expressions of genes in different cell types. **c**, Visualizations of 12 cell-type differential expression genes in the microglial.

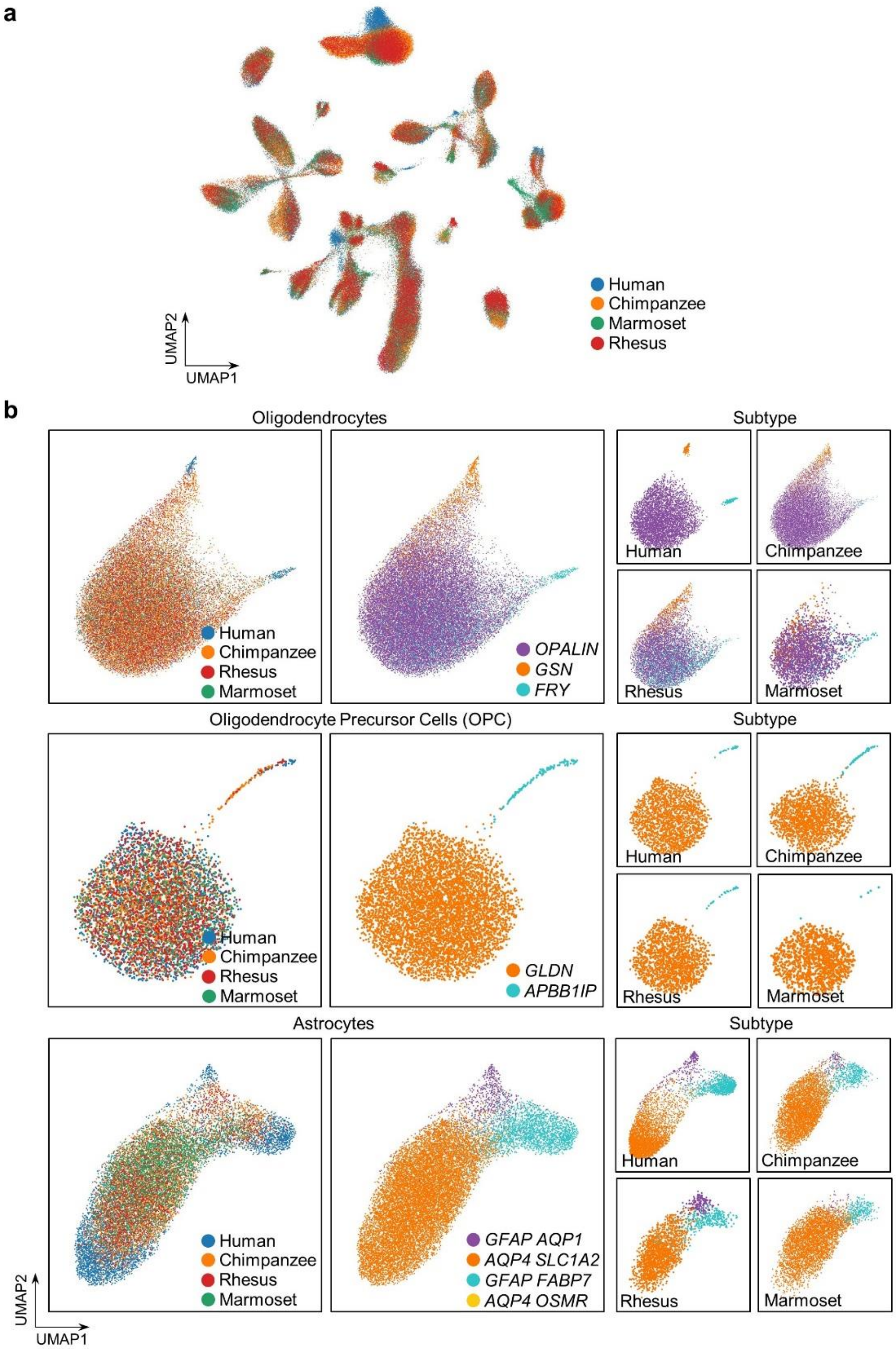


**Supplementary Figure S13.** Integrating the dorsolateral prefrontal cortex (DLPFC) dataset of four species, i.e., chimpanzees, marmosets, rhesuses, and humans by CAMEX. **a**, UMAP visualization of the whole DLPFC dataset across four species integrated by CAMEX. **b**, UMAP visualization of the astrocyte, oligodendrocyte, and oligodendrocyte precursor cell (OPC) dataset across four species mapped by CAMEX.

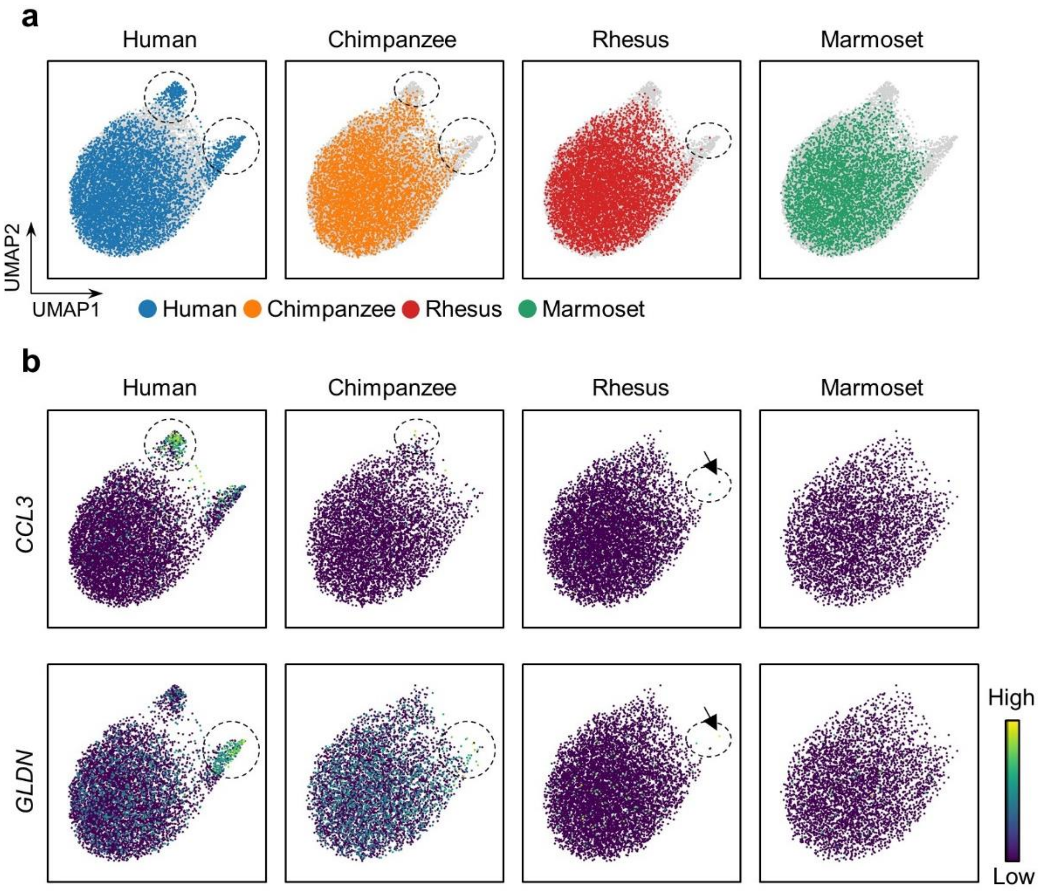


**Supplementary Figure S14.** Integrating microglia cell dataset of four species by CAMEX. **a**, UMAP visualization of the microglial dataset across four species. **b**, Visualizations of *CCL3* and *GLDN* in the microglial across four species.

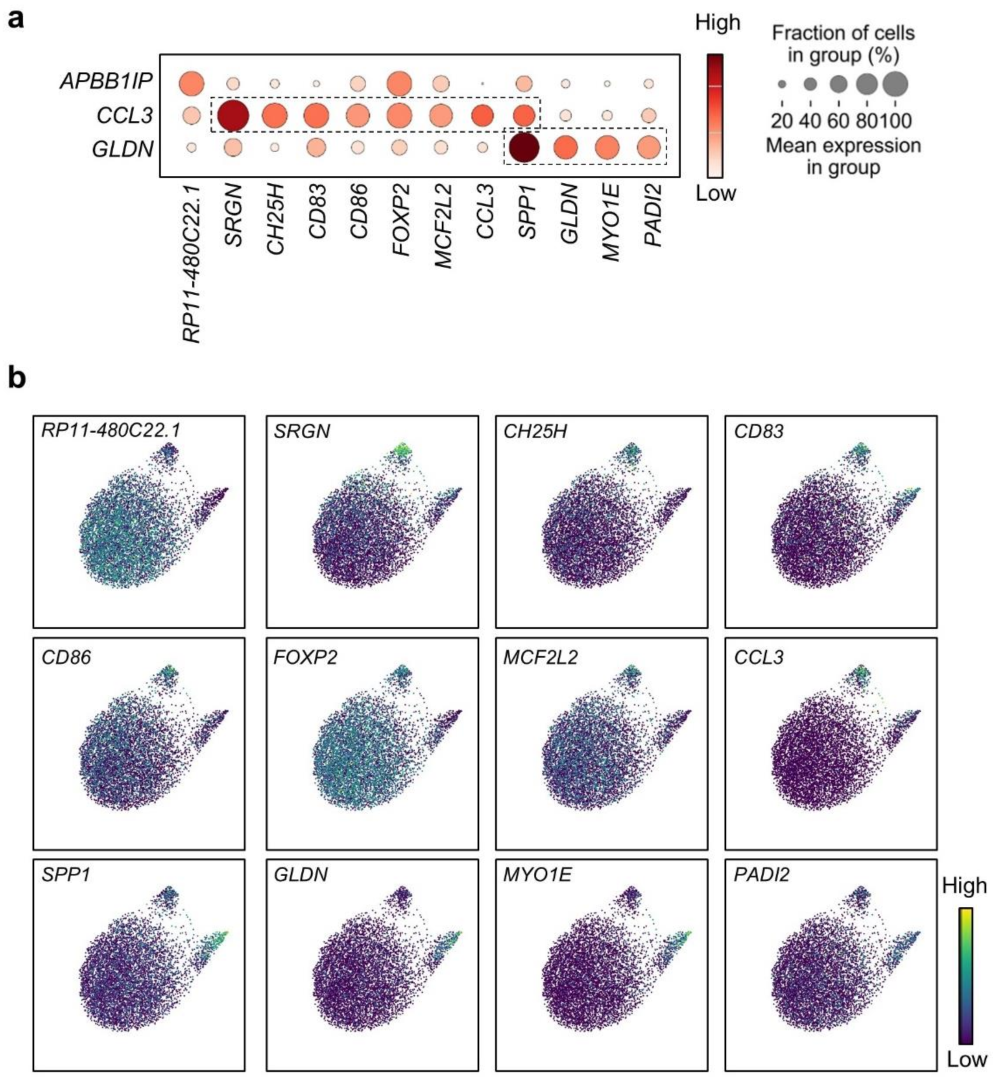


**Supplementary Figure S15.** Integrating microglia cell dataset of four species by CAMEX. **a**, Dot plots showing the expressions of genes in different cell types. **b**, Visualizations of 12 cell-type differential expression genes in the microglial.
